# *Streptococcus pneumoniae* infection of lung epithelial cells induces internalization of surface GPI-anchored proteins through pneumolysin-mediated activation of host Rho GTPases

**DOI:** 10.1101/2024.04.17.590015

**Authors:** Nick Lee, Manmeet Bhalla, Joana M. Pereira, Stacie Clark, Sara E. Roggensack, Alenka Lovy, Michael J. Pereira, Guillermo Arroyo, Walter Adams, Andrew Camilli, Rodney K. Tweten, John M. Leong, Sandra Sousa, Elsa N. Bou Ghanem

**Affiliations:** Department of Molecular Biology & Microbiology, Tufts University School of Medicine, Boston, MA; Department of Microbiology & Immunology, University at Buffalo School of Medicine, Buffalo, NY; Graduate Program in Molecular Biology & Microbiology, Tufts Graduate School of Biomedical Sciences, Boston, MA; i3S-Instituto de Investigação e Inovação em Saúde, Universidade do Porto, Porto, Portugal; Instituto de Biologia Molecular e Celular, Universidade do Porto, Porto, Portugal; Molecular and Cellular (MC) Biology PhD Program, ICBAS - Instituto de Ciência Biomédicas Abel Salazar, University of Porto, Porto, Portugal; Center for Vision Research and the Department of Ophthalmology and Visual Sciences, SUNY Upstate Medical University Syracuse, NY; Department of Biological Sciences, San Jose State University, San Jose, CA; Department of Microbiology & Immunology, The University of Oklahoma Health Sciences Center, Oklahoma City, OK; Stuart B. Levy Center for the Integrated Management of Antimicrobial Resistance, Tufts University, Boston, MA

**Keywords:** Pneumococcus, Lung Epithelial Cells, PLY, Glycosylphosphatidylinositol Anchored Proteins, GPI-AP, RhoGTPases, RhoA, Cdc42, Membrane Repair, Cholesterol-Dependent Cytolysins

## Abstract

A return to homeostasis after infection-associated cellular injury can be accelerated by a rapid damage response. *S. pneumoniae*, a typically asymptomatic colonizer of the host upper respiratory tract, can cause serious and life-threatening infections when it gains access to the lungs and other organs. The cholesterol binding *S. pneumoniae* pore-forming toxin, pneumolysin (PLY), is central to the induction of host cell damage. Here, we first found that mouse lung infection by *S. pneumoniae* diminished pulmonary expression of CD73, a glycosylphosphatidylinositol anchored protein (GPI-AP) that modulates inflammation. Infection of the human pulmonary epithelial cell line H292 resulted in a PLY-dependent reduction of not only cell surface CD73, but also the population of surface expressed GPI-APs. The decrease in cell surface GPI-APs was rapid, required pore-forming activity, and could be recapitulated by purified PLY and other cholesterol binding cytolysins. In response to PLY-mediated insult, GPI-APs were not released from the surface of epithelial cells in extracellular vesicles but rather internalized by a mechanism dependent on the Rho GTPases RhoA and Cdc42. Internalization of GPI-APs was associated with lower levels of PLY-induced apoptosis and membrane permeabilization. These findings suggest that internalization of GPI-APs from epithelial cell membranes may constitute a rapid innate repair response to cell damage induced by PLY and other pore forming toxins that could help bacteria evade host defenses as many GPI-APs have roles in immunity.

**Author summary:** *Streptococcus pneumoniae* causes serious infections that can result in mortality. The pore-forming toxin, pneumolysin (PLY) produced by these bacteria is important for their ability to cause disease. Understanding how the host responds to damage by this toxin can result in better treatment against infection. In this study, we found that PLY-mediated injury results in decreased expression of glycosylphosphatidylinositol anchored proteins (GPI-AP) from the cell surface by internalization. GPI-AP co-localize in cholesterol-rich areas of the membrane where PLY inserts to form pores and cells with decreased surface GPI-APs were associated with less of PLY-induced cell death and membrane permeabilization. These results suggest that GPI-AP are internalized as part of repair mechanisms activated in response to infection-induced cell injury. As many GPI-APs have important roles in the immune response, their removal from the cell may inadvertently help the bacteria establish better infection.

## Introduction

*Streptococcus pneumoniae* (pneumococcus), although an asymptomatic colonizer of the upper respiratory tract of healthy hosts, can, in susceptible populations, cause severe lower respiratory tract infections with high mortality and morbidity rates [1, 2]. *S. pneumoniae* predominantly causes pneumonia but can reach the bloodstream and cause serious extrapulmonary diseases, such as septicemia, meningitis, and endocarditis [3, 4]. More than 100 different *S. pneumoniae* serotypes have been identified based upon capsular polysaccharide content [5, 6], which plays a role in the severity of pneumococcal disease [7]. *S. pneumoniae* harbors many other virulence factors that promote its transition from the colonizer state to an infectious stage. These range from surface proteins involved in interaction with host pulmonary epithelium to the expression of factors that mediate metabolic adaptation or immune evasion [8].

Among these bacterial factors is a 53-kDa heat-labile cholesterol-dependent cytotoxin (CDC), pneumolysin (PLY), which is a highly conserved and important virulence factor that promotes the establishment of infection and evasion of host immune responses [9, 10]. PLY creates pores in the cell membrane [11–13] through a multistep process [14] in which 30-50 toxin monomers insert in the cholesterol-rich domains of host cells, oligomerize to form a large pre- pore complex [15, 16] that then undergoes conformational changes to produce a large 25-nm β-barrel pore in the membrane [11, 17]. Numerous studies have highlighted the role of PLY in damaging the pulmonary epithelium, activating the classical complement pathway, inducing inflammation, and inhibiting neutrophil function [8, 12]. Strains lacking PLY show reduced virulence [18].

Glycosylphosphatidylinositol-anchored proteins (GPI-APs) are expressed in highly divergent organisms, e.g., from protozoa to humans, and mediate diverse functions, such as environmental sensing, cell adhesion and migration, and immunomodulation [19–21]. GPI-APs are attached to the outer leaflet of the cell membrane through a highly conserved glycosylphosphatidylinositol (GPI) anchor, which is composed of a phospholipid portion and a glycan chain [22]. The GPI anchor is covalently linked to the C-terminus of the protein in a multi-step post-translational modification process that occurs in the endoplasmic reticulum (ER) [23, 24]. The GPI-APs then migrates to the cell surface via the secretory pathway, which involves trafficking through the ER, Golgi apparatus, and vesicular transport mechanisms, steps that are at least partially dictated by the GPI anchor [25]. Machinery that controls membrane fusion and sorting drives GPI-AP insertion in the plasma membrane, and in polarized epithelial cells, GPI-APs are selectively sorted to the apical domain [26] where they are organized into lipid rafts, specialized microdomains enriched in cholesterol and sphingolipids [22, 27]. In polarized cells the clusters are assembled in the Golgi apparatus and traffic to the apical domain, while in non-polarized cells the clusters organize from GPI-APs monomers already expressed at the cell membrane [28]. The expression of GPI-APs on the cell surface is highly regulated. GPI-APs can be released from the cell surface by the action of specific phospholipases, such as phospholipase C (PLC) [20], which cleave the GPI anchor resulting in the release of the protein into the extracellular environment. They can also be internalized through endocytosis, sorted and recycled back to the cell surface following vesicular trafficking pathways [29].

Rho GTPases are small GTP-binding proteins that act as molecular switches by cycling between an inactive GDP-bound state and an active GTP-bound state [30]. When activated, Rho GTPases interact with downstream effector proteins to initiate specific signaling cascades, which ultimately regulate key cellular functions [30]. In particular, Rho GTPases are implicated in the regulation of intracellular membrane trafficking processes such as endocytosis, exocytosis, and vesicle transport, and they control the cytoskeleton remodeling required for vesicle movement and fusion with target membranes [31]. The Rho GTPase Cdc42 has been proposed as a regulator of GPI-APs uptake [29]. During GPI-APs endocytosis, Cdc42 activation and the level of cholesterol at the cell membrane are coupled, with concomitant recruitment of actin polymerization machinery [32]. Following their internalization through a pathway that is cholesterol-sensitive and dependent on Cdc42-activation, GPI-APs traffic to the recycling endosomes and are recycled back to the cell surface, thus replenishing the pool of proteins that may have been removed or degraded.

Many GPI-APs have roles in immune modulation [20, 24]. Among them is CD73, the enzyme required for production of extracellular adenosine (EAD). EAD is produced in response to cellular damage whereby the ATP released in the extracellular milieu is sequentially converted to adenosine monophosphate (AMP) and then to adenosine by two surface ecto-nucleotidases, CD39, a transmembrane protein and the GPI-AP CD73, respectively [33]. CD73 is crucial for host resistance to infection by *S. pneumoniae* [34] as mice lacking this enzyme are highly susceptible to infection and cannot control bacterial numbers [34]. We previously reported that the absence of CD73 on neutrophils diminished their ability to kill pneumococcus [34], thus favoring bacterial growth. Interestingly, the expression of CD73 is significantly reduced on neutrophils migrating into the lungs in response to pneumococcal infection [35], indicating that pneumococcal infection may result in the modulation of CD73 on host cell surfaces.

In this study, we report that infection with *S. pneumoniae* reduces CD73 surface expression on host pulmonary epithelial cells. This reduction reflects a general effect of PLY on the surface GPI-AP expression and depends on its pore forming capacity. The reduction in CD73 expression was not due to its release from the cell surface but rather due to internalization of GPI-APs in a RhoA- and Cdc42-dependent manner. Internalization of GPI-APs was associated with reduced host cell damage in response to PLY. This study identifies a possible mechanism through which repair of damage caused by S*. pneumoniae* helps the bacteria potentially escape the host immune response.

## Results

### Infection with *S. pneumoniae* reduces the expression of GPI-APs on the surface of lung epithelial cells

We previously found that the EAD-producing enzymes CD73 and CD39 were crucial for host resistance against pneumococcal pneumonia [34, 36, 37]. To study the effect of *S. pneumoniae* infection on the expression of CD73 and CD39, we intra-tracheally (i.t.) infected C57BL/6 mice with *S. pneumoniae* TIGR4 strain or mock-treated with PBS. Forty-eight hours post infection (hpi), the lungs were harvested and the expression of CD73 and CD39 on total lung cells quantified using flow cytometry. *S. pneumoniae* significantly reduced the percentage of lung cells that express CD73 compared to uninfected controls, while the expression of CD39 remained unchanged (Fig 1A). In addition, a reduction in the amount of surface CD73 was detected on lung epithelial (EPCAM^+^) cells (Fig 1B).

**Figure 1:**
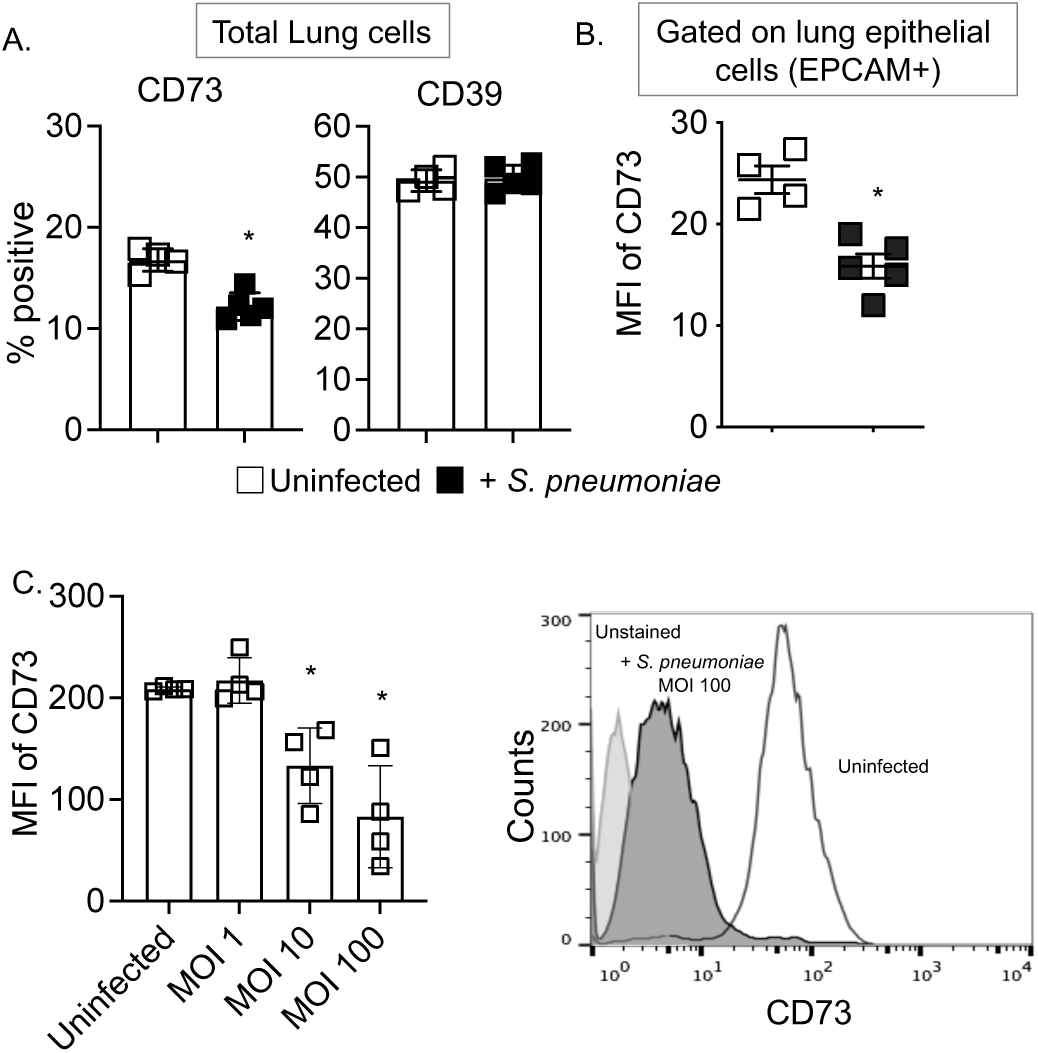
Pulmonary infection by *S. pneumoniae* significantly reduces CD73 expression on the surface of lung epithelial cells. C57BL/6 (8-weeks old) male mice were infected intra-tracheally with 2x10^6^ CFU of *S. pneumoniae* TIGR4 strain or mock infected with PBS. Mice were euthanized 48 hpi, the lungs harvested and digested into a single cell suspension, and surface stained for CD73 and CD39. Flow cytometry was used to determine (A) the percentage of CD73^+^ and CD39^+^ total lung cells and (B) the expression (mean fluorescent intensity or MFI) of CD73 on lung epithelial cells (gated on EPCAM+ cell population). The data shown are pooled from 4-5 mice per group with each box representing one mouse. *, *p*< 0.05 by Student’s t-test indicate significant differences from uninfected controls. (C) H292 cells were challenged with *S. pneumoniae* TIGR4 strain at indicated MOIs for 1 h and MFI of surface exposed CD73 was measured by flow cytometry. The panel on the right shows a representative histogram showing the intensity of CD73 labeling comparing uninfected and infected conditions. Data are pooled from four independent experiments. *, *p*< 0.05 by One-way ANOVA followed by Dunnet’s test.

To further explore the mechanisms involved, we modeled the decreased expression of CD73 on lung epithelial cells *ex vivo* using H292 cells, a human pulmonary epithelial cell line extensively used to study the interaction between the host factors and respiratory pathogens [38–41]. Infection of H292 cells with increasing numbers of *S. pneumoniae* TIGR4 resulted in a dose-dependent decrease in the expression levels of CD73 (Fig 1C), mimicking the *in vivo* finding.

These data indicate that *S. pneumoniae* infection specifically regulates the expression of CD73, a GPI-AP, but does not affect the levels of CD39, a transmembrane protein [42]. We thus tested if infection with *S. pneumoniae* selectively targets the surface expression of GPI-APs in general. H292 cells infected with *S. pneumoniae* TIGR4 were labelled with antibodies recognizing the GPI-APs CD55 and CD59 or stained with fluorescently labeled inactive variant of the protein aerolysin (FLAER), which selectively binds GPI anchors at the cell membrane [43]. Flow cytometric analysis showed that *S. pneumoniae* infection significantly reduced the expression of both CD55 and CD59, as well as FLAER staining intensity (Fig 2A). This reduction was not common to all surface proteins, as *S. pneumoniae* infection increased the expression of transmembrane proteins such as CD47 and CD54 (Fig 2B). These findings indicate that pneumococcus specifically reduces the surface expression of GPI-APs.

**Figure 2:**
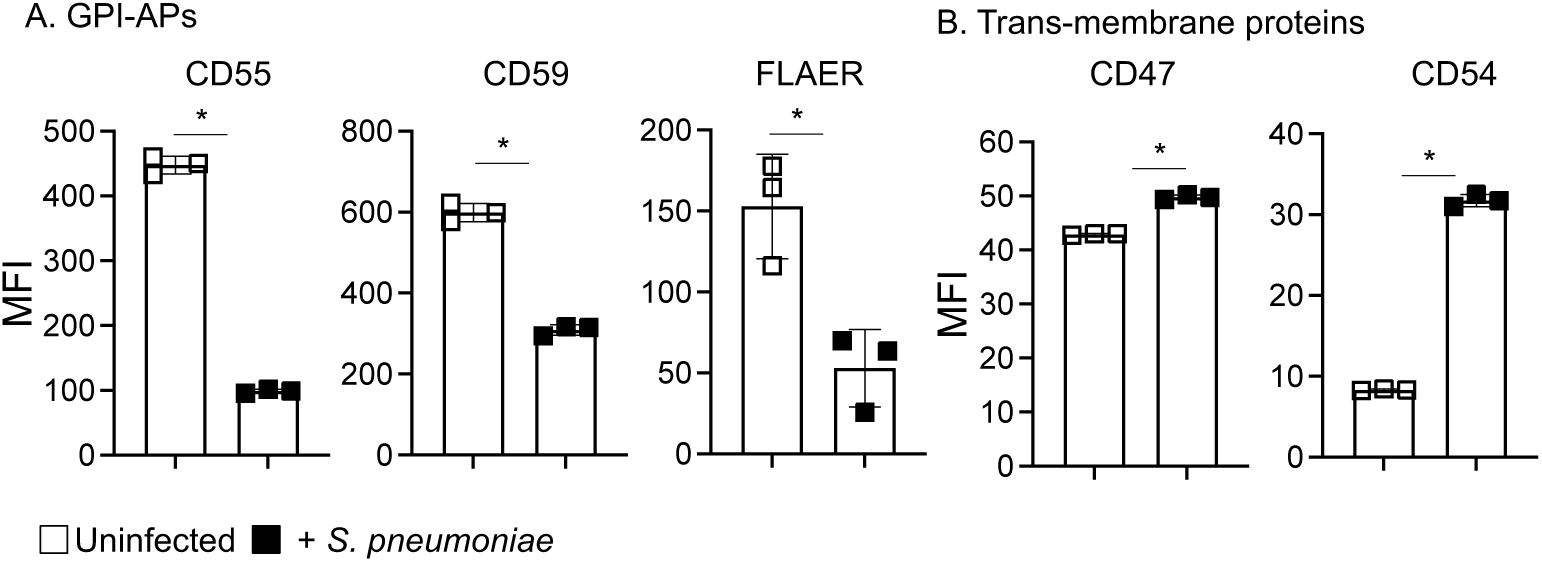
*S. pneumoniae* infection of lung epithelial cell lines decreases surface associated Glycosylphosphatidylinositol anchored proteins (GPI-APs). H292 cells were infected with *S. pneumoniae* TIGR4 strain at an MOI of 10 for 1 h. Flow cytometry was used to measure the expression (MFI) of (A) CD55 and CD59, GPI-APs detected using specific antibodies, and of bulk GPI-APs detected by using the fluorescent-labeled mutant of aerolysin (FLAER) or (B) CD47 and CD54, both transmembrane proteins detected by using specific antibodies. Data are pooled from three independent experiments. *, *p*< 0.05 by Student’s t-test indicates significant differences from uninfected controls.

*In vivo*, epithelial cells are polarized, and GPI-APs mainly localize at the apical domain [44]. To determine if the pneumococcus-induced reduction of GPI-APs surface exposure also occurs on polarized epithelial cells, H292 cells were grown as polarized monolayers on Transwell filters [39, 45]. The cells were then apically infected with *S. pneumoniae* TIGR4 for an hour and the expression of GPI-APs was analyzed through flow cytometry. The expression levels of the GPI-APs CD73, CD55 and CD59 were significantly reduced in infected cells as compared to the uninfected controls (Fig S1). Overall, these data show that infection with *S. pneumoniae* reduces the surface expression of GPI-APs on lung epithelial cells.

### *S. pneumoniae*-induced reduction of GPI-APs is dependent on the pore forming ability of PLY

To determine if the reduced surface exposure of CD73 was strain dependent, H292 were infected with D39, a serotype 2 strain of *S. pneumoniae*. Like TIGR4 infection, infection with D39 significantly reduced the expression of CD73 on H292 cells (Fig S2A), suggesting that the effect of *S. pneumoniae* on CD73 surface expression is not strain specific. Infection of H292 cells with Δ*cps* TIGR4, lacking the polysaccharide capsule, also reduced surface CD73 similarly to wild-type TIGR4 (Fig S2A), thus indicating that the reduction of CD73 expression is independent from the capsule. However, we found that heat-killed *S. pneumoniae* TIGR4 not only failed to reduce CD73 expression on H292 cells but rather increased it (Fig S2B). This indicates that a heat-labile bacterial factor is involved in reducing the expression of CD73 on lung epithelial cells.

*S. pneumoniae* pneumolysin (PLY) is heat-labile [12, 13], so we tested if the reduction in CD73 is PLY-dependent. H292 cells infected with Δ*ply* TIGR4 did not show diminished CD73 expression on the cell surface compared to the uninfected controls (Fig 3A). Moreover, infection of H292 cells with *Bacillus subtilis* expressing *ply* (*B. subtilis* PLY) significantly reduced levels of CD73 as compared to cells challenged with wild-type *B. subtilis* (Fig 3A). To test if PLY alone is sufficient for reducing the expression of CD73, H292 cells were treated with increasing concentrations of purified recombinant PLY (rPLY). The expression of CD73 was reduced upon incubation with rPLY in a dose-dependent manner (Fig 3A). Altogether, these findings demonstrate that PLY is required and sufficient for decreasing CD73 from the cell surface of pulmonary epithelial cells.

**Figure 3:**
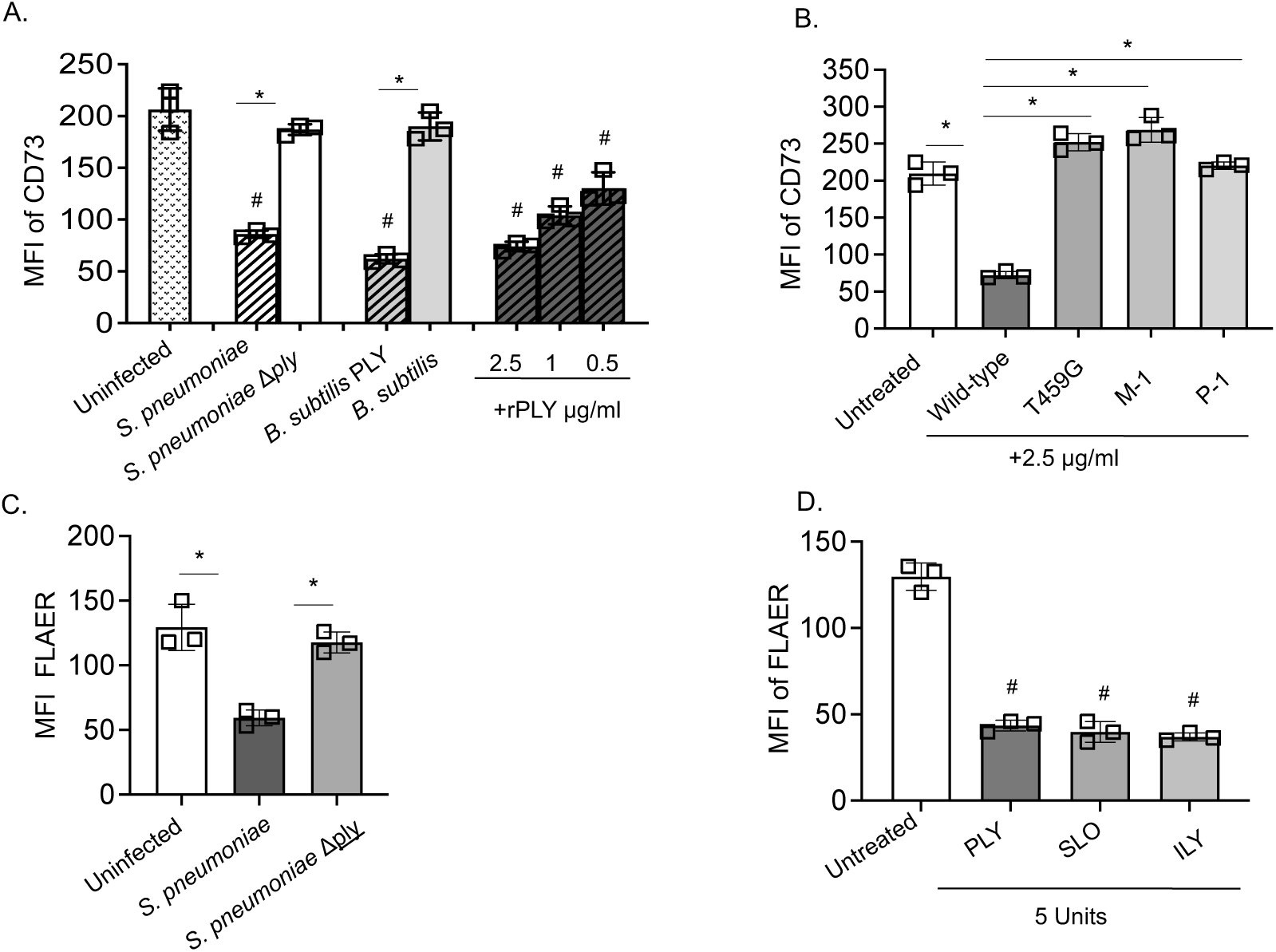
The decrease in cell surface expression of GPI-APs is dependent on PLY and its pore formation capacity. H292 cells were infected with the indicated bacterial strains at an MOI of 10 or treated with the indicated concentrations of recombinant toxins for 1h (A-B) or 10 min (C-D). The cells were then stained and the MFI of CD73 or FLAER was measured by flow cytometry. Data pooled from three independent experiments are shown. ^#^, *p*< 0.05 indicates significant differences from uninfected controls and *, *p*< 0.05 indicates significant differences between the indicated groups by One-way ANOVA followed by Tukey’s test (A-C) or Dunnet’s test (D). Units for the indicated toxins were determined by their ability to induce pores in the cells. 1U of toxin is equivalent to the concentration of toxin that causes pores in 50% of the cells. rPLY (recombinant PLY), PLY (Pneumolysin), SLO (Streptolysin O) and ILY (intermedilysin). T459G, M-1 and P-1 refer to *S. pneumoniae* strains expressing mutant forms of PLY that cannot form fully functional pores (see Materials & Methods).

To determine if pore formation is required to decrease CD73 expression on host cell surface, we tested PLY mutants defective at different stages of pore formation. 2.5 µg/ml of wild-type PLY elicited more than 3-fold reduction in CD73 expression (Fig 3A and B) but the same concentration of PLY ^T456G^, which is incapable of binding cholesterol [46], PLY ^M-1^, which is locked at an early stage of pore formation [11, 15, 47], or PLY ^P-1^, which is locked at late stage [11, 15, 47], failed to induce any change in CD73 expression on lung epithelial cells (Fig 3B). These results indicate that the phenotype observed requires the pore-forming ability of PLY.

As PLY pores may result in the death of lung epithelial cells during infection [48, 49], we tested if the reduced expression of GPI-APs could be a consequence of cell death. We tracked cell death following infection with *S. pneumoniae* TIGR4 by measuring apoptosis and found that infection resulted in rapid apoptosis of H292 cells (Annexin^+^) as early as 10 minutes post-infection (Fig S3A). This depended on PLY, as infection with Δ*ply S. pneumoniae* TIGR4 was not associated with a significant effect on cell viability (Fig S3A). However, flow cytometric analysis showed that infection with *S. pneumoniae* induced a similar significant 2-fold reduction in the intensity of FLAER on both apoptotic (Annexin^+^) and non-apoptotic (Annexin^-^) cell populations compared to the uninfected control groups (Fig S3B), demonstrating that decrease in GPI-APs is not a result of cell death. We further confirmed that PLY-dependent reduction in GPI-APs could be observed at this early 10-minute timepoint, as 10-minutes post-infection of H292 cells with wild-type but not Δ*ply* TIGR4 resulted in significant reduction of FLAER binding (Fig 3C). Altogether, these data indicate that *S. pneumoniae*-induced reduction in GPI-APs on host lung epithelial cells is PLY-mediated.

### Cholesterol-dependent-cytolysins reduce the expression of GPI-APs at the surface of lung epithelial cells

PLY belongs to a large family of cholesterol-dependent-cytolysins (CDC) that share structural and functional properties [12]. To determine if the PLY-mediated reduction of GPI-APs expression is a common feature shared among different members of CDCs, H292 cells were challenged with two other CDCs: Streptolysin O (SLO) and Intermedilysin (ILY), produced by Group A *Streptococci* and *Streptococcus intermedius*, respectively [50]. As pore formation is required for reduction of GPI-AP expression, we treated H292s at concentrations sufficient to induce equal levels of pore-formation, but that did not result in H292 cell death [11]. We found that incubation of cells with SLO and ILY with 5 units of pore forming activity caused a reduction in the FLAER intensity like that induced by PLY (Fig 3D), indicating that the decrease of GPI-AP expression is a feature common to different CDCs.

### GPI-APs are not released from the cell surface upon infection with *S. pneumoniae*

GPI-APs can be removed from the host cell surface through cleavage by phosphatidylinositol phospholipase C (PI-PLC) [51]. In fact, *S. pneumoniae* that produces the pneumococcal surface protein C (PspC) can activate PI-PLC [52]. However, we found that PspC-deficient *S. pneumoniae*, which are incapable of activating host PI-PLC, showed no defect in triggering a reduction of surface CD73 (Fig S4A). In addition, treatment of H292 cells with the phospholipase C specific inhibitor U-73122 (or its inactive analogue U-73343 [53]) did not abrogate the *S. pneumoniae*-mediated reduction in CD73 surface expression (Fig S4B). These findings indicate that reduction in cell surface associated CD73 upon *S. pneumoniae* infection does not depend on the activity of host PI-PLC.

As part of a membrane repair mechanism, host cells infected with *S. pneumoniae* can release extracellular membrane vesicles through which they expel membrane-bound bacterial products such as PLY [54, 55]. To determine if CD73 was shed in host-derived vesicles, H292 cells were treated with CellTracker Green dye CMFDA, which stains the cell cytoplasm and can be used to track host-derived vesicles. Following challenge with *S. pneumoniae* TIGR4, cell free supernatants were collected, stained with FLAER and the presence of vesicles simultaneously positive for FLAER and CMFDA was analyzed through flow cytometry. Surprisingly, we observed a significant reduction in the percentage of vesicles FLAER^+^ CMFDA^+^ released by H292 cells infected by *S. pneumoniae* compared to the uninfected controls (Fig S5), suggesting that the decrease of GPI-APs induced by *S. pneumoniae* infection is not due to the release of extracellular vesicles by the infected cells. As injured cells can also release parts of injured membranes via the formation of protrusions named blebs [56, 57], we tested the effect of treating H292 cells with blebbistatin, a drug reported to inhibit blebbing [58]. We found that treatment with blebbistatin did not affect the levels of GPI-APs (Fig S6). Collectively, these data indicate that the reduction in GPI-APs following pneumococcal infection is not due to their release from the cell surface by the activity of phospholipases, shedding in vesicles or blebs.

### Infection with *S. pneumoniae* triggers the internalization of GPI-APs

As we found that the decreased levels of GPI-APs at the cell surface are not related to their extracellular release, we tested if infection of lung epithelial cells by *S. pneumoniae* triggers their internalization. To do so, H292 cells were infected with *S. pneumoniae,* stained with FLAER to identify GPI-APs, and imaged by confocal microscopy. In conditions where H292 cells were treated with HBSS (uninfected controls) or were challenged with Δ*ply S. pneumoniae*, GPI-APs were uniformly detected on the surface of the cells (Fig 4A), following the outline of the cell membrane. However, in *S. pneumoniae* infected cells, GPI-APs lose their uniform association to the cell cortex and appear in clusters inside cells (Fig 4A, white arrows). These data suggest that infection with *S. pneumoniae* triggers the internalization of GPI-APs.

**Figure 4:**
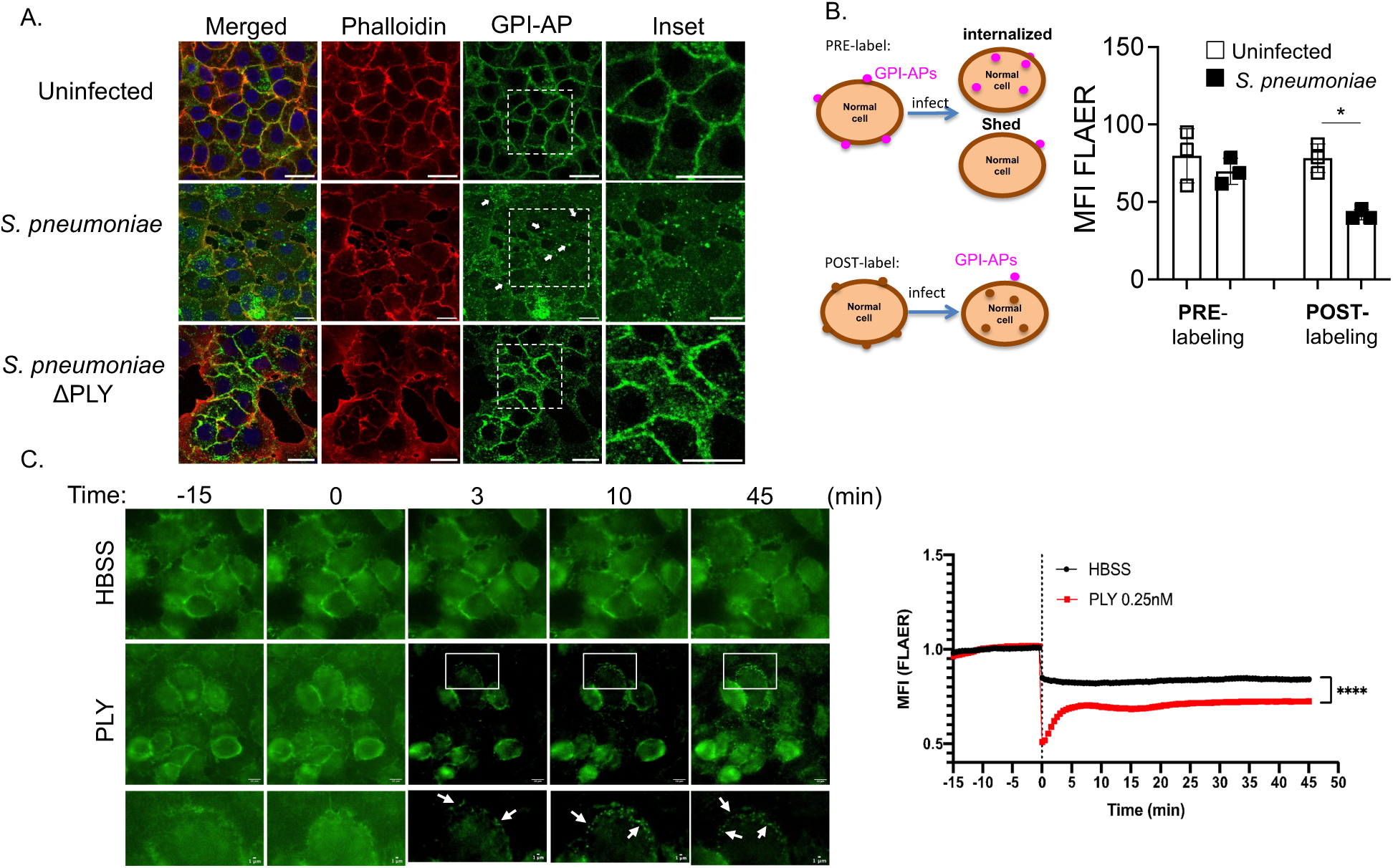
Infection with *S. pneumoniae* triggers the internalization of GPI-APs from the host cell surface. (A) H292 cells were infected with wildtype or *S. pneumoniae* Δ*ply* at an MOI of 20 for 1 h, fixed with PFA for 20 min, permeabilized with 0.1% Tx-100 and stained with phalloidin (to detect actin filaments, red) and FLAER (to detect GPI-APS, green). White arrows point GPI-APs concentrating intracellularly into vesicle-like structures. Insets highlight FLAER staining in the different conditions. Bars indicate a 20 μm scale. Representative images from four independent experiments are shown. (B) Left panel shows the schematic illustration of FLAER labeling of H292 cells either pre- or post-infection with *S. pneumoniae* TIGR4 strain (MOI of 10 for 1 h). The right panel shows the MFI of FLAER measured by flow cytometry. Data pooled from three independent experiments are shown. *, *p*< 0.05 indicates significant differences between the indicated groups by One-way ANOVA followed by Tukey’s test. (C) The left panel shows sequential frames of time-lapse microscopy video of HeLa cells pre-labeled with FLAER and challenged with PLY (0.25 nM) or HBSS (control). PLY was added at time point 0 min. The inset shows detailed distribution of GPI-APs up to 45 min post PLY challenge. The right panel shows quantified MFI for the FLAER signal overtime. Image analysis was performed using ImageJ by randomly selecting a region of interest (ROI) in the membrane of at least 53 cells per condition and calculating MFI overtime. The values were then normalized to the mean MFI calculated before adding PLY or HBSS. Scale bar for the middle lane is 10 µm and for the bottom lane is 1 µm. White arrows in the bottom lane indicate where GPI-APs appear in “vesicle-like” structures likely inside the cells. *, *p*<0.05 by Student’s paired t-test.

If GPI-APs are removed from the cell surface by internalization (vs. extracellular shedding), the total amount of GPI-AP associated with cells is predicted to be unchanged. To test that, H292 cells were stained with FLAER before infection by *S. pneumoniae* (“PRE”). After 1 h, cell-associated GPI-APs was quantitated by flow cytometry. Indeed, we found that FLAER staining remained unchanged, i.e., was indistinguishable from FLAER staining of uninfected cells (Fig. 4B, “PRE”). To confirm that under these experimental conditions, *S. pneumoniae* resulted in the removal of GPI-APs from the cell surface, in parallel we stained H292 only after the 1 h infection (“POST”). As predicted, surface staining by FLAER after infection revealed decreased surface GPI-APs (Fig. 4B, “POST”). These findings confirm that infection with *S. pneumoniae* induces the internalization of GPI-APs.

To test whether PLY was sufficient to trigger this process, as well as to follow GPI-AP internalization in living cells, HeLa cells (chosen due to their ease of imaging by microscopy) were pre-labelled with FLAER and then challenged with purified PLY or mock-treated. FLAER-stained GPI-APs were visualized by fluorescent microscopy for 45 minutes (Supplementary video 1). FLAER intensity was rapidly reduced on the cell surface upon challenged with PLY (Fig 4C and Supplementary video 1), in addition FLAER signal was detected intracellularly within 3 minutes of incubation with PLY (Fig 4C and Supplementary video 1) in structures resembling intracellular vesicles (Fig 4C and Fig S7). At 45 min after PLY incubation FLAER signal at the surface of the cells appeared to increase (Figure 4C and S7), suggesting that internalized GPI-APs are recycled back to the cell surface. Overall, these data show that the infection with *S. pneumoniae* triggers the internalization of surface GPI-APs which are then found intracellularly in vesicle-like structures.

### Internalization of surface GPI-APs is mediated by PLY-induced activation of host Rho GTPases RhoA and Cdc42

To explore the mechanism behind *S. pneumoniae*-mediated internalization of GPI-APs, we first assessed a possible role of the actin cytoskeleton, given that dynamic changes in the actin machinery are often required for endocytosis [59]. H292 cells were pre-treated with Cytochalasin D, an inhibitor of actin polymerization, at 2 μM, a concentration that prevented the actin-dependent cell-cell spread of *Listeria monocytogenes* ([60] and data not shown). Treatment with Cytochalasin D did not affect the reduction of FLAER intensity following *S. pneumoniae* infection (Fig S8A), suggesting an actin-independent mechanism for the internalization of GPI-APs. Similar results were observed when Cytochalasin D pre-treated H292 cells were challenged with PLY or SLO (Fig S8B), indicating that other CDCs also induce the internalization of GPI-APs through a mechanism independent of actin polymerization.

The host Rho family of guanosine 5’-triphosphate (GTP)-binding proteins (GTPases) are known to play important roles in protein trafficking, receptor-mediated endocytosis, and internalization of GPI-APs [61]. To examine the role of Rho GTPases in the internalization of GPI-APs induced by *S. pneumoniae* infection, H292 cells were treated either with *Clostridium difficile* toxin B (ToxB), which irreversibly inactivates Rho GTPases by glucosylation [62], or with vehicle control, before infection with *S. pneumoniae*. While infection with *S. pneumoniae* in vehicle-treated cells caused the expected reduction in FLAER intensity indicating the decreased surface expression of GPI-APs (Fig 5A), cell surface expression of GPI-APs was not affected in cells pre-treated with ToxB (Fig 5A), suggesting a role for Rho GTPases in PLY-mediated GPI-APs internalization. To confirm the role of Rho GTPases in the internalization of GPI-APs, we knocked down their expression using RNAi. H292 cells were transfected with siRNAs targeting Cdc42, RhoA and Rac-1 or with scrambled non-targeting siRNA as control. At 48 h post transfection, cells were challenged with *S. pneumoniae* at a MOI of 10 for 1 h and FLAER intensity was quantified through flow cytometry. Depletion of Cdc42 and RhoA but not Rac-1 abrogated the expected reduction in FLAER intensity usually observed in H292 cells infected by *S. pneumoniae* (Fig 5B). This indicates that the internalization of pneumococcal-mediated GPI-APs depends on the Rho GTPases Cdc42 and RhoA.

**Figure 5:**
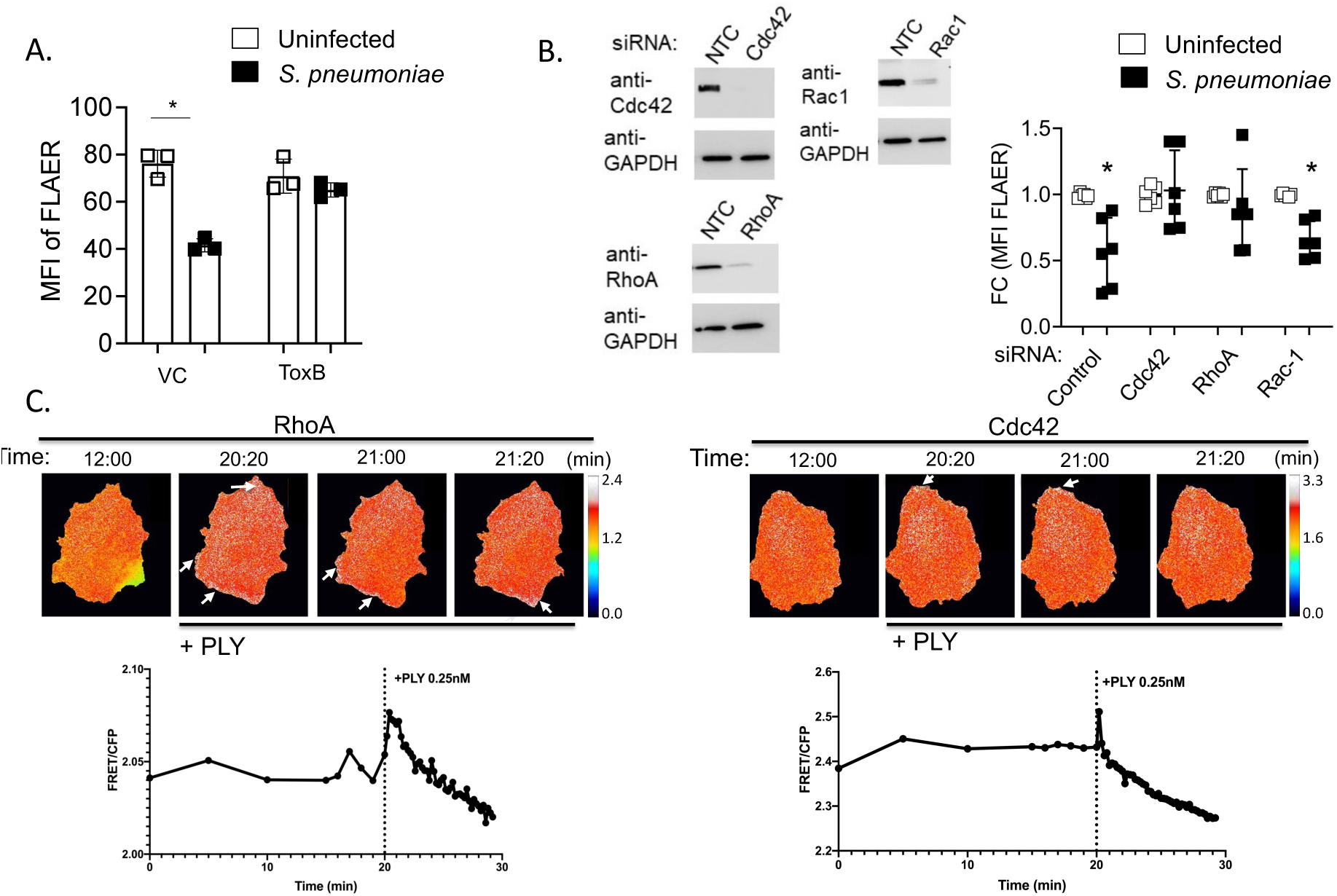
Removal of GPI-APs from the cell surface largely depends on host GTPases RhoA and Cdc42 activity. (A) H292 cells were pre-treated with 100 ng/ml *C. difficile* toxin B (Tox B) or vehicle control (VC) overnight and infected with *S. pneumoniae* TIGR4 strain at an MOI of 10 for 30 min, labeled with FLAER and the MFI was assessed by flow cytometry. Data are pooled from three independent experiments. *, *p*< 0.05 by One-way ANOVA followed by Tukey’s test. (B) H292 cells were transfected with a control non-targeting (NTC) siRNA, or siRNA targeting GTPases Cdc42, RhoA and Rac1. At 48 h post-transfection, whole cell lysates were used to confirm the depletion of the target proteins through Western blotting. Detection of GAPDH was used as a loading control (left panel). Depleted and control cells were infected with *S. pneumoniae* TIGR4 at an MOI of 10 for 30 min, labeled with FLAER and the MFI assessed by flow cytometry (right panel). The MFI values of FLAER are displayed as fold changes with respect to the uninfected baseline for the respective siRNA condition. The data shown are the mean of three independent experiments done in duplicates. *, *p* < 0.05 indicates significantly different than 1 by one-sample t-test. (C) HeLa cells over-expressing a FRET probe for RhoA or Cdc42 were imaged for 20 min before and 10 min after incubation with 0.25 nM of PLY. The upper panel shows representative sequential frames of time-lapse images of HeLa cells before and after incubation with PLY. FRET/CFP ratio images are shown in blue-to-white pseudocolor scale with white being high activation. Arrows show the regions where RhoA (left) or Cdc42 (right) are activated (FRET/CFP signal is high). The lower panel shows the quantitative analysis for the ratio between the FRET and CFP signal (FRET/CFP), which measures the RhoA and Cdc42 activation levels. The quantification was performed using a custom-made macro to calculate FRET/CFP ratio specifically at the cell membrane. 79 cells (for RhoA) and 74 cells (for Cdc42) were analyzed from two independent experiments.

Rho GTPases were previously reported to be activated by PLY in neuronal cells [13, 63]. We assessed if Cdc42 and RhoA are activated by PLY in epithelial cells in the time frame of PLY-mediated internalization of GPI-APs. We transfected HeLa cells with plasmids encoding the fluorescence resonance energy transfer (FRET) probes (CFP/YFP FRET pairs) that reflect the activation state of RhoA or Cdc42. Each probe comprises a truncated form of the GTPase and the Rho-binding domain (RBD) of effectors, bound to the CFP or YFP, respectively. The binding of GTP to the GTPase (resulting in activation) brings the YFP-bound GTPase domain and the CFP-bound RBD into close proximity, allowing FRET from CFP to YFP [55, 56]. Transfected HeLa cells were challenged with 0.25 nM of PLY and live-cell microscopy was performed to visualize the activation status of RhoA and Cdc42. FRET ratio (FRET/CFP) was calculated specifically at the cell cortex using Image J. Cells were imaged for 20 min before PLY was added and for additional 10 min after PLY challenge. For both probes, the ratio of FRET/CFP remained stable during the 20 min before PLY addition, showing that the levels of GTPase activation do not vary in these conditions (Fig 5C). Shortly after the addition of PLY, we observed a modest but consistent increase in the FRET/CFP ratio levels. The increase detected for the Cdc42 probe was very rapid and of short duration (about 40 s), while the increase for the RhoA probe appeared more sustained in time (140s) (Fig 5C). These data suggest that PLY triggers a fast and transient activation of both RhoA (Supplementary video 2) and Cdc42 (Supplementary video 3) at the cell cortex. The ratio of FRET/CFP, for both probes, rapidly decreased to values that tended to be lower than those obtained before PLY incubation (Fig 5C), indicating that although the activity of RhoA and Cdc42 is triggered during the very early response to PLY induced damage, it is rapidly shutdown. Overall, these data indicate that PLY induces internalization of GPI-APs through the rapid, spatially, and temporally regulated activation of the Rho GTPases RhoA and Cdc42.

### GPI-APs are not required for PLY-mediated pore formation

The PLY-related CDC ILY utilizes the human GPI-AP CD59 as a receptor to induce oligomerization and pore formation [64] and the cytolytic pore forming toxin aerolysin binds the GPI moiety of the GPI-anchored T-lymphocyte protein Thy-1 [19]. To test whether PLY requires the expression of GPI-APs on the host cell surface to form pores, H292 cells were either treated with recombinant PI-PLC to cleave GPI-APs from the cell surface [51], or mock-treated with PBS. As expected, treatment of cells with PI-PLC caused a significant reduction in the percentage of GPI^+^ cells (FLAER^+^) compared to the untreated control group (Fig S9A). Upon addition of PLY to H292 cells, we detected the expected decrease in GPI^+^ cells compared to the untreated control group (Fig S9A). We did not see a further reduction in the number of cells expressing GPI-APs when PLY was added to PI-PLC pre-treated cells (Fig S9A).

We then assessed whether reduced expression of GPI-APs impact PLY-mediated pore formation using PI permeability assays. We observed that PLY challenge induced a 4-fold increase in the number of permeabilized cells (PI^+^) in both the untreated and PI-PLC pre-treated conditions in comparison to the respective baseline controls (Fig S9B). Finally, the average intensity (MFI) of PI staining was unaltered by PI-PLC treatment. These data indicate that PLY does not require surface GPI-APs to bind and form pores on host cell membranes.

### Internalization of GPI-APs is associated with lower levels of PLY-induced cellular damage

PLY forms pores in cholesterol rich domains, where GPI-APs are also colocalized [12]. Depending on the extent of damage, host cells can repair the damaged membrane and revert to homeostasis [65]. To couple the kinetics of the PLY-mediated removal of GPI-AP from the cell surface with an analysis of the population of cells that have undergone this reduction, H292 cells were challenged with *S. pneumoniae* or mock-treated with PBS for different time periods and the kinetics of reduction in FLAER intensity was determined throughout infection. According to our previous live-cell imaging experiments of cells challenged with rPLY (Fig 4C), we expected the reduction to occur rapidly. Indeed, *S. pneumoniae* significantly reduced FLAER intensity within 10 minutes of infection compared to the PBS-treated cells and retained this reduced level for the remainder of the 90-minute observation period (Fig 6A). To analyze the specific population of cells that reduced GPI-AP, we utilized flow cytometry and found that by 10 minutes after infection approximately 60% of the infected cells showed reduced levels of GPI-AP (termed “GPI^low^” cells) (Fig 6B), a percentage that also remained stable through the rest of the experiment.

**Figure 6:**
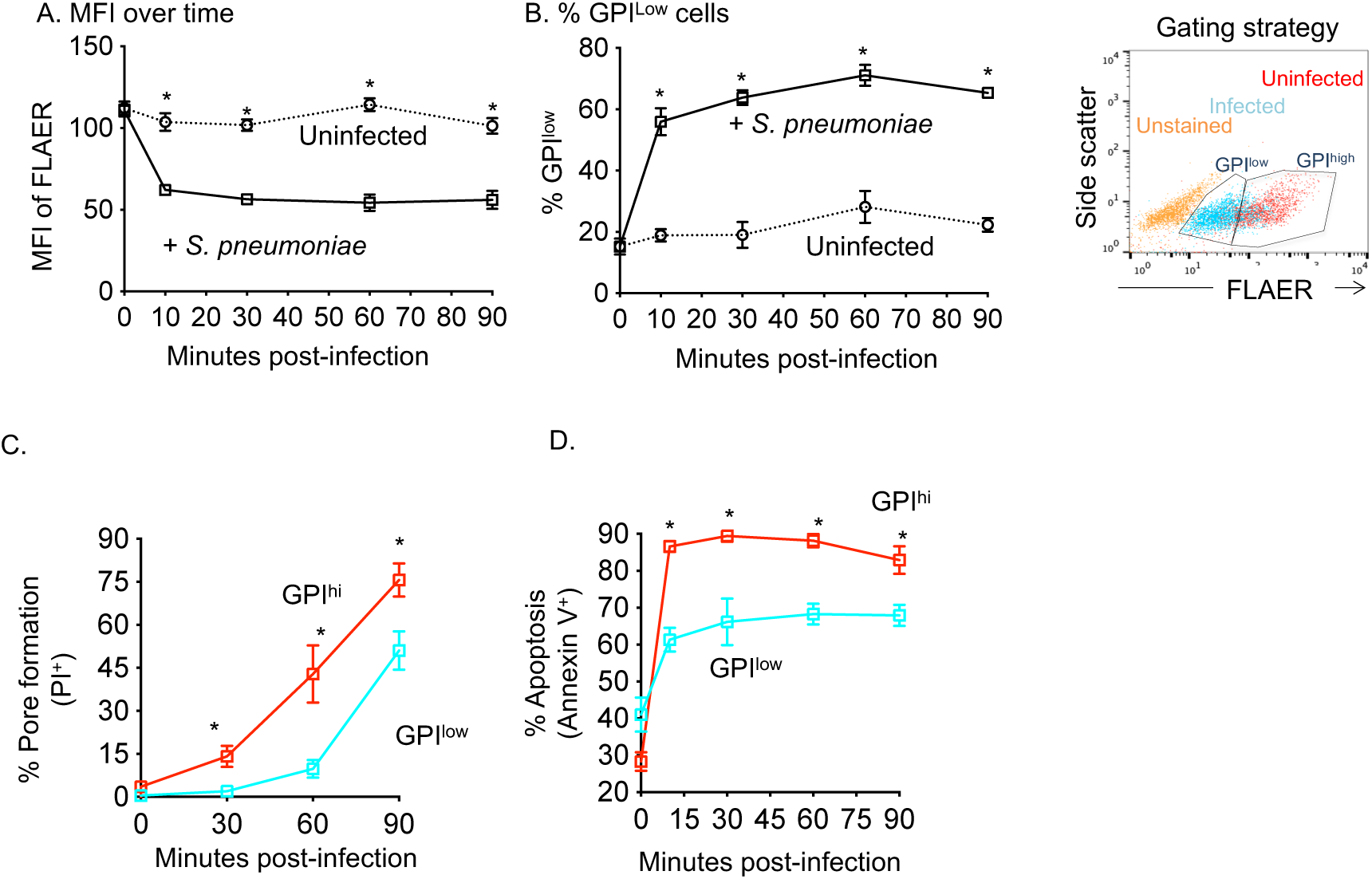
Internalization of GPI-APs is associated with lower levels of PLY-induced cellular damage. H292 cells were infected with *S. pneumoniae* TIGR4 strain at an MOI of 10 or left uninfected. The cells were then labeled with FLAER and flow cytometry was used to measure (A) the MFI of FLAER at indicated time points of infection and (B) calculate the percentage of H292 cells (left panel) with down-regulated FLAER expression (termed as GPI^low^) based upon the gating strategy shown in the right panel. (C) H292 cells suspended in media containing propidium iodide (PI) were infected with *S. pneumoniae* TIGR4 strain at an MOI of 10 and labeled with FLAER. GPI^low^ vs GPI^hi^ cell populations were then gated on, as indicated in the left panel of (B) and the percentage of H292 cells within each population that incorporated the PI dye was assessed by flow cytometry. (D) H292 cells were infected with *S. pneumoniae* for the indicated times and stained with Annexin V and FLAER. GPI^low^ vs GPI^hi^ populations were gated (right panel in B) and the percentage of apoptotic (Annexin V +) cells within each population was determined through flow cytometry. Data pooled from three independent experiments are shown. * *p*< 0.05 indicate differences by one-way ANOVA comparison between the different groups at indicated time points.

To test if the reduction of GPI-APs from the cell surface affected cellular viability, we measured the staining of the population of GPI^low^ or GPI^high^ H292 cells by Annexin V, which stains apoptotic cells, and PI, which stains cells with permeable membranes. Interestingly, we found that, throughout infection, the population of infected GPI^high^ H292 cells had a significantly higher percentage of membrane damaged (PI^+^) (Fig 6C) and apoptotic (Annexin V^+^) cells than the population of GPI^low^ cells (Fig 6D). These findings suggest that reduced GPI-APs surface expression is associated with reduced levels of PLY-induced damage. Altogether, these findings suggest that cells with reduced expression of GPI-APs are more resistant to PLY-induced toxicity and that the internalization of GPI-anchored proteins seems to be part of membrane repair that limits the extent of PLY-induced cellular damage.

## Discussion

The outcome of host-pathogen interaction is proposed to be mediated by the damage response [66, 67]. This damage can be caused by the invading organism and/ or the host’s immune response to infection. *S. pneumoniae*, a typically asymptomatic colonizer of the host upper respiratory tract, can cause serious and life-threatening infections when it gains access to the lungs and other organs and this is directly linked to its ability to cause organ damage and impair organ function [68–71]. One key component of this is the bacterial toxin PLY, found in all the clinical isolates of *S. pneumoniae* [72]. PLY induces host cell damage by making pores in cells [71]. However, host survival can be contingent on repairing/controlling infection-induced damage [73]. In this study, we found that PLY-induced pores are associated with the rapid removal of GPI-APs from the host cell surface, which can have profound effects on host response to infection. PLY activated host RhoGTPases RhoA and Cdc42, resulting in the internalization of GPI-APs (Figure 7). This internalization of GPI-APs was associated with reduced cellular damage thus, suggesting the induction of host repair mechanisms.

**Figure 7:**
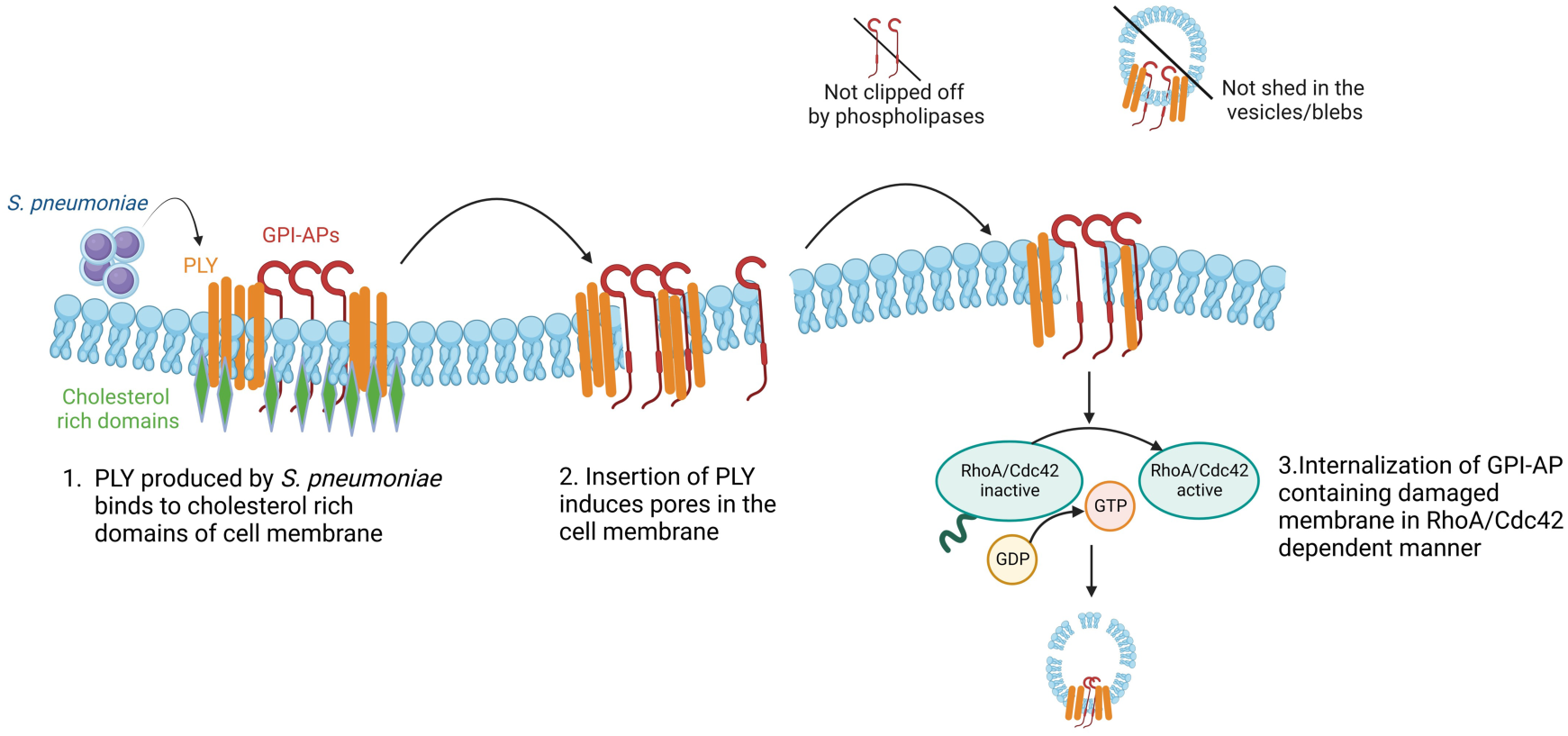
Working Model. PLY inserts in cholesterol-rich microdomains where GPI-APs are colocalized. PLY-induced pores trigger membrane repair resulting in internalization of the GPI-AP containing damaged membrane in a RhoA/Cdc42-dependent manner.

GPI-APs are a heterogenous group of proteins which lack transmembrane domains and are dependent on the GPI moiety for their expression on the cell surface [24]. Through their interactions with the sphingolipids and cholesterol in the lipid bilayer, the GPI moieties anchor the proteins in lipid-rich microdomains or membrane rafts [24, 74]. We found here that removal of GPI-APs from the cell surface was not only mediated by PLY, but also occurred upon treatment of host cells with other cholesterol dependent cytolysins (CDCs) including SLO and ILY. CDCs bind to the cholesterol moieties in the host cell surface through the undecapeptide domain of the peptide to mediate their cytolytic activity [14, 64]. Binding of the soluble monomeric form of the CDCs to the cholesterol-rich membrane results in a sequence of structural changes causing oligomerization of the monomers with subsequent formation of transmembrane β-barrel that results in pore formation in the host cell membrane [64, 75]. Therefore, CDCs and GPI-APs will likely localize together in cholesterol-rich microdomains at the cell surface where the damage occurs. Consequently, internalization of GPI-APs following CDC-induced damage may be part of membrane repair. Prior work with SLO showed that pores formed by the toxin are endocytosed as part of the membrane repair process [76]. Here, we found that removal of GPI-APs by PLY was dependent on the pore forming ability of the toxin and that internalization of GPI-APs was associated with reduced cellular damage, suggesting that the host can repair the damage at the membrane.

The type and extent of host cell damage as well as the host response to CDCs is highly concentration dependent [65]. At high concentration levels, PLY induces irreversible loss of cell membrane integrity and cellular homeostasis, while at lower concentrations, it activates host cell repair mechanisms to restore their membrane integrity and to survive [65]. Initiation of host cell repair of the plasma membrane depends on a limited influx of Ca^2+^ that triggers cytoskeletal rearrangements to plug the holes in the plasma membrane [13, 77]. This has been described to be mediated by the activation of host GTPases, re-localization of host intracellular organelles and recruitment of cytoplasmic Ca^2+^ responsive proteins including annexins that interact with the plasma membrane phospholipids at the injury site [13, 77, 78]. Host Rho GTPases, including Rho, Rac and Cdc42, play a major role as molecular switches that regulate key cellular signaling pathways and link surface receptors to the downstream processes culminating in endocytosis, exocytosis and actin polymerization that affect cell polarity, migration, cytokinesis and vesicle trafficking [79]. While exploring the cellular mechanisms that drive GPI-APs reduction at the cell surface, we found that removal of GPI-APs from the cell surface upon *S. pneumoniae* infection was dependent on Cdc42 and RhoA while Rac1 was not involved in this process. These data fit with what has been reported in the literature regarding the role of Rho GTPases in the turnover of GPI-APs under homeostatic conditions. GPI-APs are normally recycled from the cell surface through an endocytic process called potocytosis involving caveolae, which is independent of both dynamin and clathrin-mediated pathways [29, 80]. Once internalized, GPI-APs within the cell can be seen in tubular structures called GPI-enriched endosomal compartments [80, 81]. Internalization of GPI-APs via endocytosis is under the regulation of host GTPases [80, 81]. This includes CD59, one of the GPI-APs for which we observed reduction in surface expression upon infection, whose internalization via endosomes is reported to be regulated by Cdc42 [82]. Additionally, perturbation in cholesterol moieties in the lipid bilayer also leads to internalization of GPI-APs via clathrin-and dynamin-independent endocytic pathway [80]. Therefore, CDC binding to the cell surface and pore formation results in the perturbation of the cholesterol rich microdomains where GPI-APs are localized resulting in initiation of the endocytic pathway which may lead to the repair of the injured membrane.

One of the interesting findings of this study was that actin polymerization had no apparent role in the internalization of GPI-APs despite the activation of Cdc42 and RhoA. Cdc42 is a key regulator of actin polymerization through the regulation of N-WASP and Arp2/3 complex and similarly, RhoA also regulates actin polymerization [79]. Several recent reports have described actin-independent endocytic events in various cell types. Work on non-adherent K562 human erythroleukemic cells or adherent Cos-7 cells showed a lack of actin polymerization role in the endocytosis process in these cells [83]. In another study, treatment of A431 cells (adherent and in suspension) with cytochalasin D did not affect the process of endocytosis indicating an actin-independent mechanism of endocytosis in these cells [83]. Additionally, the rational that the polymerization of actin is important to generate the mechanical force required for vesicle fission during clathrin-mediated endocytosis was recently challenged. In mouse chromaffin cells, inhibition of dynamin GTPase but not of actin polymerization resulted in changes in the fission-pore conductance in the endocytic membranes indicating that the pinching off of the endocytic vesicle from the plasma membrane is actin-independent [84]. The requirement of actin in endocytosis could be spatially regulated as in polarized cells inhibition of actin polymerization blocked clathrin-mediated endocytosis from the apical but not the basolateral surface of polarized Madin-Darby Canine Kidney Cells (MDCK) [85]. This study observed PLY-mediated reduction in GPI-AP expression in pulmonary epithelial cells from infected murine models as well as in non-polarized and polarized H292 cells. Overall, our findings point to a signaling pathway whereby PLY activated Cdc42 and RhoA initiate internalization of GPI-APs independent of actin polymerization machinery. Other bacterial pathogens have evolved to use actin polymerization-independent mechanisms to trigger host responses. For example, *Salmonella enterica* serovar Typhimurium was reported to activate Rho kinases to mediate entry into the cell by activating myosin II independent of actin polymerization [86]. Similarly, *Listeria monocytogenes* was shown to manipulate host exocytosis independent of actin polymerization to enter the host cell [87].

Apart from stimulating membrane repair through annexins, injury to plasma membrane by PLY, LLO or SLO also results in a phenomenon called shedding whereby the host cell releases extracellular vesicles to not only remove the injured part of the membrane but also to remove the membrane bound toxin [54, 88]. Blebbing is one of the ways to mediate the shedding of larger vesicular structures whereby the cell undergoes actomyosin-dependent localized contraction in the plasma membrane to create confined spaces or a localized bulge with elevated Ca^2+^ levels to prevent leakage to cytoplasmic contents and proteins in the extracellular milieu [13, 65]. These localized bulges in the plasma membrane are eventually released by the cells either to remove the areas of pores in case of a low degree of plasma membrane injury or to initiate the processes of regulated apoptosis and inflammation in case of extensive cell damage [56, 65]. Since shedding and blebbing result in the loss of part of the plasma membrane, it could also result in the loss of expression of GPI-APs. However, data presented here using CMFDA dye to track the vesicles and using chemical inhibition of blebbing showed that this was not the case. Surprisingly, we observed a significant reduction in the percentage of vesicles FLAER+ CMFDA+ released by H292 cells infected by *S. pneumoniae* compared to the uninfected controls. We also found no evidence for the enzymatic release of GPI-APs from the cell surface to the extracellular milieu. Certain bacterial species including *S. pneumoniae* have evolved to induce the expression of host lipolytic enzyme phospholipase [89]. Given that the anchors of GPIs interact with the lipid moieties of the plasma membrane, the GPI-APs are therefore susceptible to cleavage by phospholipases [20, 21]. The findings here with phospholipase C specific inhibitor U-73122 and using a Δ*pspC S. pneumoniae* strain showed that the reduction in CD73 expression following PLY treatment was not a consequence of cleavage of GPI-APs by the host phospholipase.

The removal of GPI-APs from the host cell surface can seriously affect the host’s ability to control infection. These GPI-bound proteins are essential for many host physiological functions including but not limited to enzymatic activities, nutrient uptake, complement regulators, immune antigens and adhesion molecules [20, 28, 34, 90, 91]. This work shows that treatment with PLY reduces CD55, CD59 and CD73 levels on pulmonary epithelial cells, all of which are GPI-APs and important regulators of host inflammatory response. CD73 is an ectonucleosidase that is required for initiating host responses to damage [42]. The host can sense infection-induced damage by recognizing several damage-associated molecular patterns (DAMPS), including direct recognition of extracellular ATP and its breakdown product adenosine in the extracellular environment [92]. CD73 is required to dephosphorylate ATP, which has leaked from damaged cells, into adenosine [93]. We previously found that extracellular adenosine production by CD73 was required for host resistance against *S. pneumoniae* in part by modulating epithelial responses to these bacteria [34–36]. Other groups have also found that CD73 expression on epithelial cells is important in other infections where CD73 expression on intestinal epithelial cells was needed to control *Salmonella enterica* serovar Typhimurium infection [94]. Apart from CD73, both CD55 and CD59 act as inhibitors of complement membrane attack complex (MAC) [95, 96] and have a role in neutrophil trans-epithelial migration [97]. Therefore, PLY’s downregulation of CD55 and CD59 could also have important implications on host immune responses. In summary, our findings suggest that repair mechanisms activated in response to infection-induced cellular damage may inadvertently help *S. pneumoniae* and other bacterial pathogens evade host defenses, reflecting the dual function of PLY during infection.

## Materials and Methods

### Ethics Statement

Mice were handled in accordance with the recommended Guide for the Care and Use of Laboratory Animals published by the National Institutes of Health. All procedures were done according to the protocol approved by the University at Buffalo Institutional Animal Care and Use Committee (IACUC), approval number MIC33018Y.

### Mice

All the animal work was performed in 8-12 weeks C57BL/6J male mice purchased from Jackson Laboratories (Bar Harbor, ME) and housed at a specific-pathogen free facility at the University at Buffalo.

### Mammalian cell lines and media

Human epithelial cervical cancer cell line HeLa (ATTC) and pulmonary mucoepidermoid carcinoma cell line NCL-H292 (ATCC) were grown in Dulbecco’s modified Eagle’s medium (ThermoFisher Scientific) and RPMI 1640 media (ThermoFisher Scientific), respectively, supplemented with 10% fetal bovine serum (FBS) (ThermoFisher Scientific) and 100 U penicillin/streptomycin (ThermoFisher Scientific). For all the experiments involving lifting and/or seeding, the cells were washed twice with dPBS (Gibco) and treated with Trypsin/EDTA (Gibco).

### Bacterial strains and growth conditions

The following bacterial strains were used: *S. pneumoniae* TIGR4 and D39 [98], PLY-deletion (Δ*ply*) [98], capsule-deletion (Δ*cps*) [99], and pneumococcal surface protein C-deletion (Δ*pspC*) mutant strains of *S. pneumoniae* TIGR4 [100] *B. subtilis* and its mutant expressing PLY (*B. subtilis* PLY) [101]. The pneumococcal strains were grown to mid-log phase in Todd–Hewitt broth (BD Biosciences) supplemented with Oxyrase (Fisher) and 0.5 % yeast extract at 37 °C/5 % carbon dioxide. Bacterial aliquots were then frozen at –80 °C in growth media with 20 % glycerol. Tryptic Soy Agar plates with 5 % sheep blood (Northeast Laboratory Services, P1100) were used to confirm the titers of *S. pneumoniae* in the aliquots. Before infecting animals or H292 cells, bacterial aliquots were thawed and diluted in PBS to the desired number. *B. subtilis* strains were grown overnight at 37 °C in Luria-Bertani (LB) Broth.

### Toxin purification

Purification of SLO (Streptolysin O), ILY (intermedilysin) and PLY (PLY) along with its mutant forms was carried out as previously described [11, 47]. Briefly, the cultures of *E. coli* BL-XL-1 Blue expressing polyhistidine-tagged toxins were grown in sterile TB broth (with 1:100 of overnight culture) grown at 37 °C with shaking at 200 rpm supplemented with 100 μg/mL ampicillin. Expression of toxins was induced by the addition of isopropyl β-D-thiogalactopyranoside (IPTG, Fisher Scientific) to a final concentration of 1 mM when the OD600 of the culture reached 0.5-0.6 after which the cultures were allowed to grow overnight. This was followed by centrifugation, resuspending and lysing bacterial pellets in PBS supplemented with halt protease inhibitor cocktail (Thermo Fisher Scientific) using microfluidizer. The cell lysates were centrifuged at 10000 g for 30 min at 4 °C and the clear supernatant was collected and passed through Ni-NTA agarose column (Qiagen). Contaminating proteins were washed away by passing a 20–120 mM imidazole gradient through the column while the bound toxins were eluted using PBS containing 500 mM imidazole. The purity of the toxin was confirmed by running the collected fraction on SDS-PAGE. The toxin-containing fractions were mixed with 10% (v/v) glycerol and 5μM Tris(2-carboxyethyl) phosphine hydrochloride (TCEP) and flash-frozen on dry ice and then shifted to −80 °C until use. *C. difficile* toxin B (Tox B) was purchased from Sigma. Unless indicated, we used 0.25 nM of toxin for experiments with purified PLY as this concentration corresponded to 1 pore forming Unit which causes PI uptake in only 50% of the cell population and was sufficient to induce low-grade cellular damage.

### Toxin activity assay

The activity of purified toxins was measured as previously described [11]. Briefly, aliquots of different toxins were thawed on ice and centrifuged at 14,000 rpm at 4 °C for 10 min to remove protein precipitates. Bradford assay (Bio-Rad) was used to measure the concentrations of toxins per the manufacturer’s instructions. Toxins at different concentrations were added in wells of non-adherent 96-well plates containing approximately 1 x 10^5^ H292 cells and incubated for 10 min at 37 °C. Following incubation, cells were stained with propidium iodide (PI) (1 mg/mL), incubated in dark at RT for 15 min and analyzed through flow cytometry. The toxin concentration that caused permeabilization of 50 % of the H292 cells (PI+) was referred as “1 unit” (1U) of toxin activity. Cells treated with 5 % Triton X-100 acted as a positive control while HBSS was used as negative control.

### Murine infection studies

Mice anesthetized under isoflurane treatment were intratracheally (it) instilled with *S. pneumoniae* TIGR4 at a dose of 2 x 10^6^ CFU, as described previously [34, 102]. At 48 h post-infection, the mice were euthanized, lungs were harvested and single-cell suspension was obtained by digesting the tissue in RPMI 1640 containing 10 % FBS and 2 mg/mL of type 2 collagenase (Worthington) and 30 µL/mL of 10 mg/mL DNase (Worthington) stock prepared in dH2O as previously described [102].

### Preparing heat killed *S. pneumoniae* TIGR4

Frozen aliquots of *S. pneumoniae* TIGR4 were thawed on ice, centrifuged at 1700 g for 5 min and resuspended in PBS in the original volume. The bacterial suspension was then incubated at 65 °C for 2.5 h. Bacterial killing was confirmed by plating the suspension on blood agar plate.

### Preparation of polarized H292 monolayers and infection with *S. pneumoniae*

Polarized H292 monolayers were prepared as previously described [45]. Briefly, the inverted side of the filter membrane of 24-well Transwells with a pore size of 3 µm were coated with 70µL of 30µg/mL of sterile collagen solution (3 mg/mL of collagen stock diluted in 60% ethanol) and allowed to dry for 3 to 4 h at RT under sterile conditions. Approximately 1 x 10^6^ H292 cells resuspended in RPMI 1640 media (with FBS and Pen/Strep) were added in the volume of 70µL on the coated filter membrane and allowed to grow overnight at 37 °C with 5 % CO2. The following day, the Transwells with cell were inverted into a sterile 24-well plate containing 1mL of H292 complete media and cells were allowed to grow and polarize for a week at 37 °C with 5 % CO2. Cells were then washed, infected apically with *S. pneumonia,* treated with trypsin and type II collagenase, stained for GPI-APs, and analyzed by flow cytometry.

### Antibodies, inhibitors, and other reagents

The antibodies used in flow cytometry studies to measure the expression of specific GPI-APs included the following antibodies: anti-CD73 (AD2, eBioscience), anti-CD39 (eBioA1, eBioscience), anti-CD55 (JS11, Biolegend), anti-CD47 (CC2C6, Biolegend) and anti-CD54 (HA58, Biolegend). The primary antibodies used for Western blotting included anti-Rac1 (Proteintech) (1:1000), anti-Cdc42 (Cell Signaling) (1:1000), anti-RhoA (Cell Signaling) (1:1000) and anti-GAPDH (MA5-15738) (1:5000). Horseradish peroxidase (HRP)-conjugated secondary antibodies were purchased from Jackson Immunolabs. The following anti-mouse antibodies were also used: Anti-CD326 (EpCAM) (G8.8, eBioscience), anti-CD39 (24DMS1, eBioscience) and anti-CD73 (eBioTY/11.8, eBioscience). The inhibitors Cytochalasin D, PI-PLC, PI-PLC inhibitor U-73122 and its control U-73343 (Invitrogen, ThermoFisher Scientific) and Blebbistatin were purchased from Sigma and dissolved in DMSO (unless indicated). As appropriate, control groups were treated with diluted DMSO, PBS or HBSS.

### Detection of plasma membrane integrity

*S. pneumoniae* or PLY-induced cellular damage was determined by using ANNEXIN/PI staining (BioLegend), per the manufacturer’s protocol. Briefly, infected H292 cells were centrifuged at 1200 rpm at 4 °C, the supernatants were discarded, and pellets were re-suspended in 100 µL of Ca^2+^ free staining buffer (supplied with the kit). 2.5 µL of FITC Annexin V and 5 µL of Propidium Iodide were used per sample. The samples were then incubated for 15 min at room temperature (RT) in the dark and immediately analyzed by flow cytometry.

### FLAER labeling

Following challenge with different bacterial strains or toxins, H292 cells were centrifuged at 1200 rpm at 4 °C, the supernatants were discarded, and pellets were re-suspended in 50 µL of FLAER (FL2S, Cederlane) diluted in FACS buffer (HBSS Ca/Mg^+^ containing 1% FBS) at the final dilution of 1:100. The cells were then incubated for 30 min on ice in the dark. Following incubation, the cells were washed twice with 100 µL of FACS buffer, resuspended in 300 µL of FACS buffer and analyzed by flow cytometry.

### Flow cytometry

Fluorescence intensities for different GPI-APs, Annexin V and Propidium Iodide were measured on a BD Fortessa cytometer. At least 50,000 events for lung tissue and 10,000 events for *ex vivo* assays with H292 cells were captured for analysis. For the analysis of FLAER-positive cell vesicles (Figure S5), the events were captured by legacy MoFlo cytometer. For the data in Figure S9, the BD Accuri C6 flow cytometer was used to capture the events. All the analysis was done using FlowJo software.

### RNA-interference

Approximately, 1.5 x 10^4^ H292 cells were seeded in 24-well plates (Falcon) and cells were transfected the following day in serum free media using 100 nM of short interfering RNA (siRNA), Opti-mem (Invitrogen) and the lipid reagent LF2000 (Invitrogen) as described previously [1]. Serum was added on day 3 at a final concentration of 10% in the transfected wells. On day 4, the cells were challenged with *S. pneumoniae* TIGR4 strain or lysed for Western blotting to confirm protein depletion. SMARTpool siRNAs from Dharmacon were used to deplete Cdc42 (M-005057-01-0005), Rac1 (M-003560-06-0005) and RhoA (M-003860-03-0005). A non-targeting NTC siRNA (D-001210-02-05) was used as a control.

### Western blotting

To confirm protein depletion, approximately 48 h post-transfection H292 cells were lysed on ice for 30 min in ice-cold RIPA buffer (1% Triton X-100, 0.25% sodium deoxycholate, 0.05% SDS, 50 mM Tris-HCl [pH 7.5], 2 mM EDTA, 150 mM NaCl, 1 mM phenylmethylsulfonyl fluoride, 1 mM sodium orthovanadate, 10 mg/liter each of aprotinin and leupeptin), as previously described [103, 104]. The cell lysates were centrifuged at 12,000 rpm at 4 °C and bicinchoninic acid kit (Pierce) was used to determine the protein concentration in the samples. An equal amount of protein (20 µg) was run on Mini-PROTEAN TGX Stain-Free Precast Gels (BioRad) SDS-polyacrylamide gel and transferred to polyvinylidene difluoride (PVDF) membranes. The membranes were blocked in 3% BSA for 1h at RT and incubated overnight at 4 °C with primary antibodies diluted in TBS/T with 3 % BSA. The membranes were washed thrice with TBST (5 min per wash) followed by incubation with HRP-tagged secondary antibodies for 1 h at RT. The bands were detected using ClarityTM Western ECL substrate (BioRad) on the ChemiDoc XRS+ system (BioRad).

### Measuring phosphatidylinositol-specific phospholipase C (PI-PLC) and PLY-induced cell damage

H292 cells were seeded in a 6-well plate (5 x10^5^ cells/well) and 20 h later were pre-treated with PI-PLC (0.1U) prepared in (Opti-MEM supplemented with 10mM Hepes, 1mM EDTA and 0.1%BSA) [105]. Cells were then challenged for 10 min with PLY at 0.25 nM diluted in HBSS (or HBSS only in the untreated condition). Cells were then harvested, centrifuged at 1200 rpm for 5 min at 4 °C and incubated for 30 min on ice in 50 µL of FACS buffer containing 0.5 µL of FLAER (1:100). Following incubation, cells were washed and centrifuged at 1200 rpm for 5 min at 4° C and resuspended in FACS buffer. Just before flow cytometry analysis, propidium iodide (PI) was added at 2 ug/mL and incubated for 1 minute. Then, cells were analyzed for FLAER and PI staining in BD Accuri C6 flow cytometer. At least 20,000 events per sample were collected. Data obtained were analyzed using FlowJo software (Version 10.8).

### Immunofluorescence microscopy

Approximately, 2 x 10^5^ H292 cells were grown in a complete medium on 13 mm diameter coverslips in 6-well plates. The day after, the cells were washed and challenged with *S. pneumoniae* in serum-free RPMI 1640 medium. The plate was centrifuged at 1000 rpm for 2 min to enhance contact between bacteria and H292 cells and then incubated for 1 h at 37 °C in 5 % CO2. Cells were washed in PBS, fixed in PBS with 4 % paraformaldehyde, quenched with 20 mM NH4Cl (20 min), permeabilized with 0.1 % Triton X-100 (5 min) and blocked with 3 % BSA in PBS (1 h). The antibodies were then diluted in PBS containing 1% BSA and cells were labeled for GPI-AP nucleus and F-actin. GPI-AP using FLAER-Alexa 488 (Cedarlane) (1:50), DAPI (Sigma) (1:1000) and Alexa fluor 680 phalloidin (Life technologies) (1:50), respectively as described previously [106]. Cells were then washed and mounted with Vectashield (Vector laboratories) and imaged using a 63 x 1.4 NA objective lens on a confocal laser-scanning microscope (Leica SP8) and processed using ImageJ software.

### RhoA and Cdc42 activation and FRET

HeLa cells seeded in Ibitreat 8-well dishes (4 x 10^4^ cells/well) (Ibidi) were transfected approximately 20 h later with 300 ng of plasmid encoding FRET probes (CFP/YFP FRET pairs) for RhoA or Cdc42 using jetPrime transfection reagent (Polyplus) and incubated for 16–20 h to allow the expression of the recombinant proteins. FRET probes used were described in [107, 108]. Cells were then kept in Opti-MEM (Gibco, without phenol red) at 37 ℃ in 5 % CO2 and imaged using Nikon TI microscope coupled with IRIS9 camera and using 60 x 1.4 NA objective. Cell imaging was started 20 min before PLY (0.25 nM) addition and continued for 10 min after challenge with the toxin. CFP/FRET/YFP fluorescence data were acquired after every 20 s. Image sequences were analyzed using ImageJ and FRET/CFP signal ratio were calculated only at the cell cortex using a custom-made macro. A total number of 79 cells from 2 independent experiments were analyzed for RhoA activation and 74 cells were analyzed from 2 independent experiments for Cdc42 activation.

### Live cell imaging and signal quantification

HeLa cells were seeded in 8-well Ibidi plates (1x10^5^ cells/well) and 20 h later stained with FLAER (1:100) for 30 min. Cells were then imaged using a Nikon Ti microscope coupled with IRIS9 camera with 40x 0.95 NA objective, for 15 min with a time interval of 30 s before the addition of PLY (0.25 nM diluted in HBSS) or HBSS (used as untreated control), and then for additional 45 min post PLY addition. Quantification of the FLAER signal was obtained by image analysis using ImageJ. A total of 53 cells were analyzed. The mean fluorescence intensity (MFI) was determined overtime in randomly selected regions of interest (ROIs) at the cell membranes. These values were then normalized to the mean MFI obtained from the same cells before these were challenged with PLY or HBSS.

### Statistical analysis

Graph Pad Prism9 was used to perform all the statistical analysis. All the data graphs represent the mean values +/- standard deviation (SD). Significant differences were determined using One-way ANOVA followed by Tukey’s or Dunnett’s multiple comparison test or Student’s t-test or one sample t and Wilcoxon test, as indicated. *p* values of 0.05 or lower were considered significant.

## Acknowledgements

We want to thank Mafalda Sousa (ALM facility, i3S) for developing the macro for calculation of the FRET/CFP signal ratio.

## Author contributions

NL and MB conducted research, analyzed data, and wrote paper. JMP, SER, AL, MP, GA, and WA conducted research and analyzed data. AK and RKT provided valuable reagents and feedback. SS and JML designed the research and wrote the paper. ENBG designed research, performed experiments, and wrote the paper. JML, SS and ENBG had responsibility for the final content. All authors read and approved the final manuscript.

