## Supplemental Figures for "*Streptococcus pneumoniae* infection of lung epithelial cells induces internalization of surface GPI-anchored proteins through pneumolysin-mediated activation of host Rho GTPases"

**Supplemental Data**

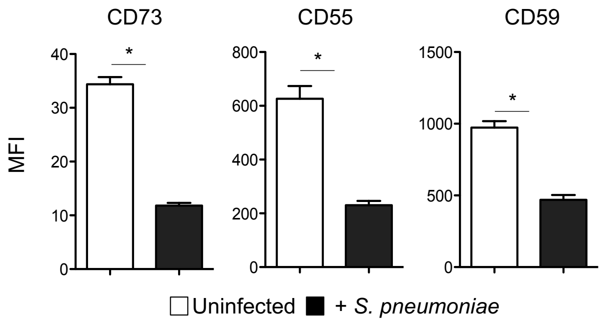


**Figure S1: *S. pneumoniae* reduces the expression of GPI-APs on polarized cells*.*** H292 cells seeded on transwells were allowed to polarize for 1-week. The cells were then infected apically with *S. pneumoniae* TIGR4 strain at an MOI of 10 for 1 h, stained for the indicated proteins and MFI was measured by flow cytometry. Representative data from one of three independent biological replicates are shown where each condition was tested in triplicate (n = 3 technical replicates per experiment). *, $p < 0.05$ by Student's t-test indicates significant differences from uninfected controls.

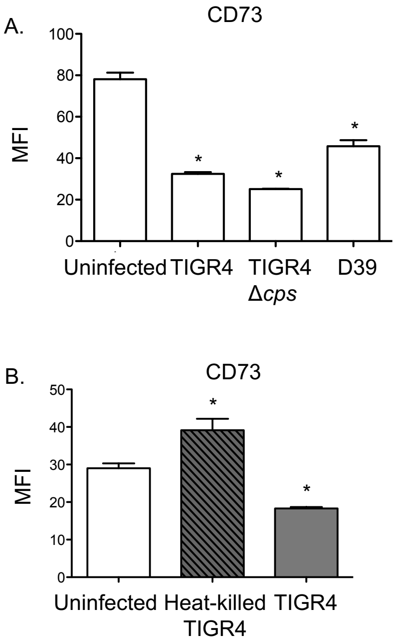


**Figure S2: *S. pneumoniae*-induced reduction in CD73 expression is not strain- or capsule-dependent but mediated by a heat-labile bacterial factor**. (A-B) H292 cells were infected with the indicated strains of *S. pneumoniae* at an MOI of 10 for 1 h and the expression of CD73 (MFI) measured by flow cytometry. Representative data from one of three independent experiments are shown where each condition was tested in triplicate (n = 3 technical replicates per experiment). *, $p < 0.05$ by one-way ANOVA comparison with uninfected controls.

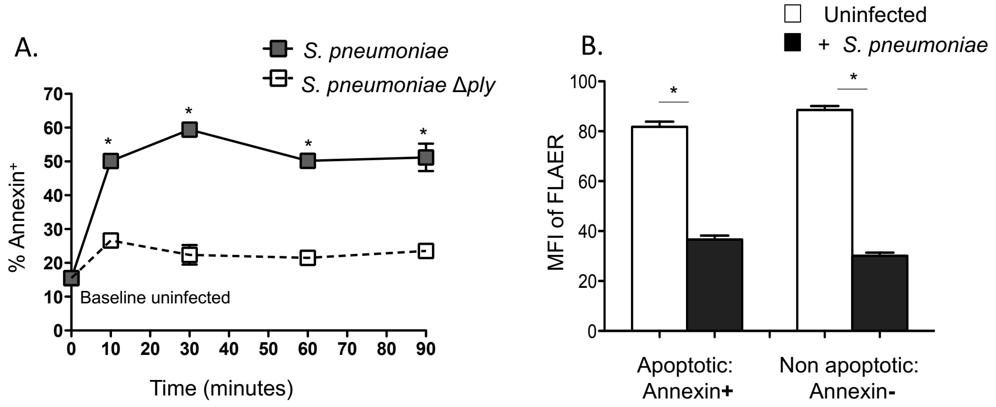


**Figure S3: PLY-induced decrease in GPI-APs expression does not result from cell death.** H292 cells were infected with wildtype or *S. pneumoniae* Δ*ply* at an MOI of 10. The cells were then stained with Annexin V and FLAER and analyzed by flow cytometry to determine (A) the percentage of apoptotic (Annexin V+) cells throughout infection and (B) the MFI of FLAER of both apoptotic (gated on Annexin V+ cells) and non-apoptotic (gated on Annexin V- cells) cell populations at the 10 min timepoint. Representative data from one of three independent experiments where each condition was tested in triplicate (n = 3 technical replicates per experiment) are shown. *, $p < 0.05$ by one-way ANOVA comparison with corresponding time points (A) and (B) uninfected controls.

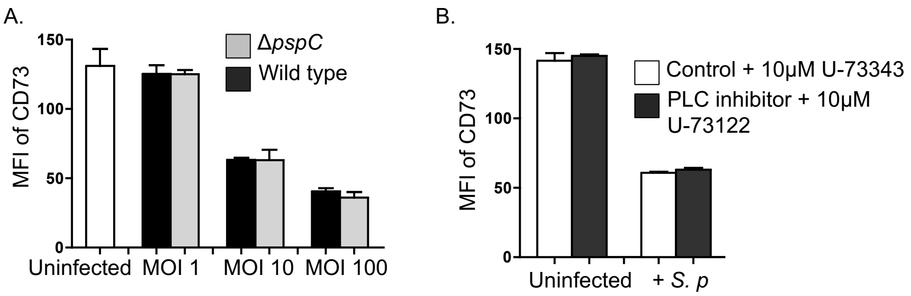


**Figure S4: PLY-induced decrease of surface CD73 expression is not mediated by host phospholipases (PLC).** (A) H292 cells were infected with wildtype or *S. pneumoniae* Δ*pspC* at indicated MOIs for 1 h and analyzed for CD73 expression through flow cytometry. (B) H292 cells were treated with 10 μM PLC inhibitor U-73122 or its inactive form U-73343 for 30 min followed by infection (in the presence of inhibitors) with *S. pneumoniae* TIGR4 strain at an MOI of 10 for 1 h. Cells were then stained and the expression (MFI) of CD73 was measured by flow cytometry. Representative data from one of three independent experiments where each condition was tested in triplicate (n = 3 technical replicates per experiment) are shown.

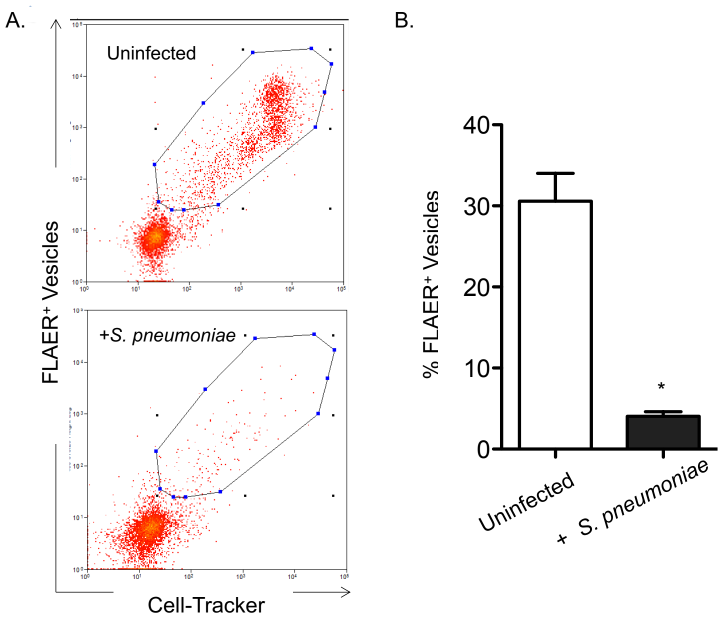


**Figure S5: *S. pneumoniae* infection decreases the release of vesicles containing GPI-APs from lung epithelial cells.** H292 cells were loaded with 1 μM of CellTracker Green dye CMFDA and challenged with *S. pneumoniae* TIGR4 strain at an MOI of 10 or left uninfected for 1 h. The cell culture supernatants were collected, centrifuged and stained with FLAER. The presence of FLAER-positive vesicles (under CMFDA+ gate) was analyzed by Legacy MoFlo cell sorter. (A) shows a representative dot plot of vesicles analyzed and (B) shows the quantification of samples under each condition. Data shown are representative from one of three independent experiments where each condition was tested in triplicate (n = 3 technical replicates per experiment). *, $p < 0.05$ by Student's t-test, indicate significant difference from the uninfected controls. CMFDA (5-chloromethylfluorescein diacetate).

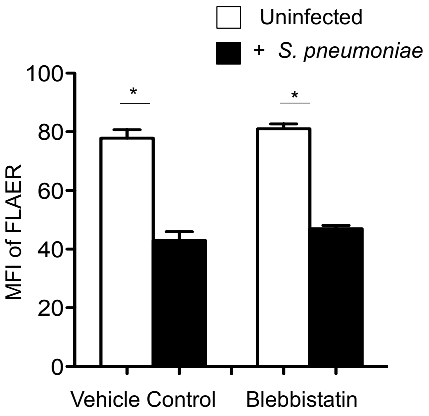


**Figure S6: *S. pneumoniae*-mediated decrease in GPI-APs expression is independent of blebbing.** H292 cells were pre-treated with 50 μM Blebbistatin or vehicle control for 20 min followed by infection with *S. pneumoniae* TIGR4 strain at an MOI of 10 for 1 h. The cells were then stained with FLAER and MFI analyzed by flow cytometry. Data shown are representative from one of two independent experiments where each condition was tested in triplicate (n=3 technical replicates per experiment). *, $p < 0.05$ by one-way ANOVA comparison with uninfected controls.

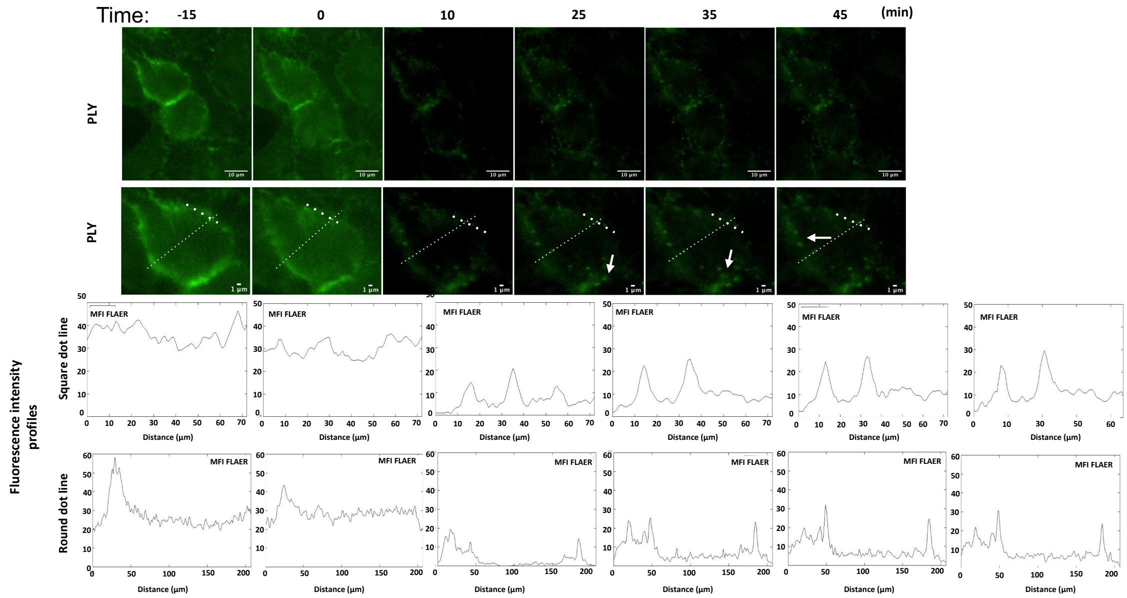


**Figure S7**: **GPI-APs are detected inside the cells in structures resembling intracellular vesicles upon PLY challenge.** Sequential frames of time-lapse microscopy video of HeLa cells pre-labeled with FLAER. PLY (0.25 nM) was added at time point 0 min. The inset (bottom row) shows details of GPI-APs distribution. FLAER intensity in line scans (lower panels) was determined along the dotted lines drawn in the inset images. Fluorescence intensity profiles (indicating FLAER distribution) at the bottom were obtained in a cortical region (thick line, upper panel profiles) and across the entire cell (thin line, lower panel profiles). Scale bar for the top lane is 10 µm and for the bottom lane is 1 µm.

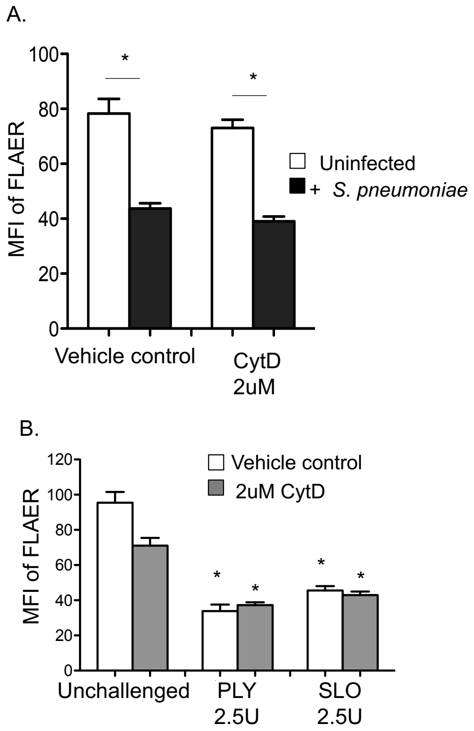


**Figure S8: PLY-mediated decrease in GPI-APs surface expression is not dependent on actin polymerization.** H292 cells were pre-treated with 2 µM cytochalasin D (CytD) or vehicle control for 30 min followed by challenge with (A) *S. pneumoniae* TIGR4 strain at an MOI of 10 for 30 min or (B) 2.5 U of recombinant PLY or SLO for 15 min. Cells were then labeled with FLAER and the MFI was assessed by flow cytometry. The graphs show representative data from one of three independent experiments where each condition was tested in triplicate (n = 3 technical replicates per experiment). *, $p < 0.05$ indicates significant differences by one-way ANOVA comparison with corresponding uninfected controls under the same treatment conditions.

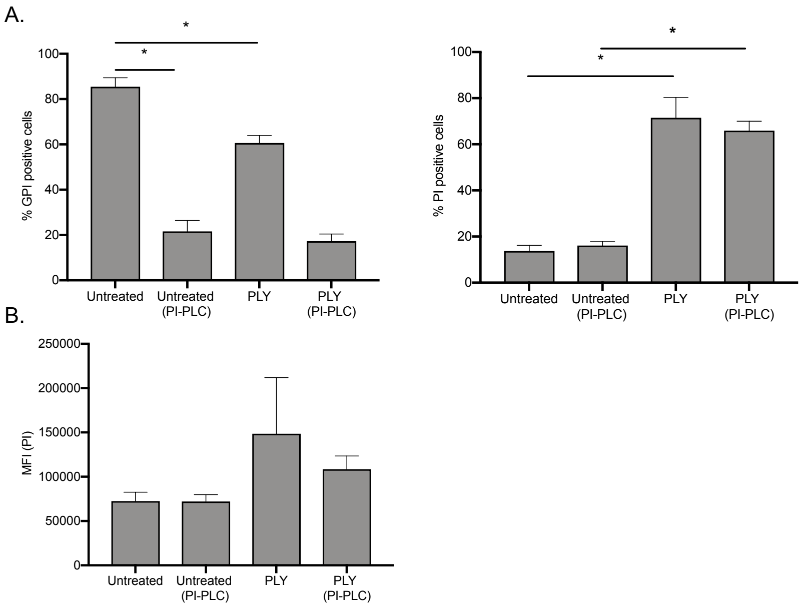


**Figure S9: Surface-exposed GPI-APs are not required for PLY-induced plasma membrane permeabilization.** (A) H292 cells incubated with or without PI-PLC (0.1 U, 2 h at 37 ℃) were challenged with purified PLY (0.25 nM) for 10 min or left untreated. Cells were then collected and processed for flow cytometry analysis to determine the percentage of GPI$^+$ and PI$^+$ cells. Data shown are pooled from at least three independent experiments. *, $p < 0.05$ by one-way ANOVA comparison with untreated cells. (B) The mean fluorescence intensity (MFI) of the PI is shown in (B).
