## Supplemental Videos for "*Streptococcus pneumoniae* infection of lung epithelial cells induces internalization of surface GPI-anchored proteins through pneumolysin-mediated activation of host Rho GTPases"

### Slide 1
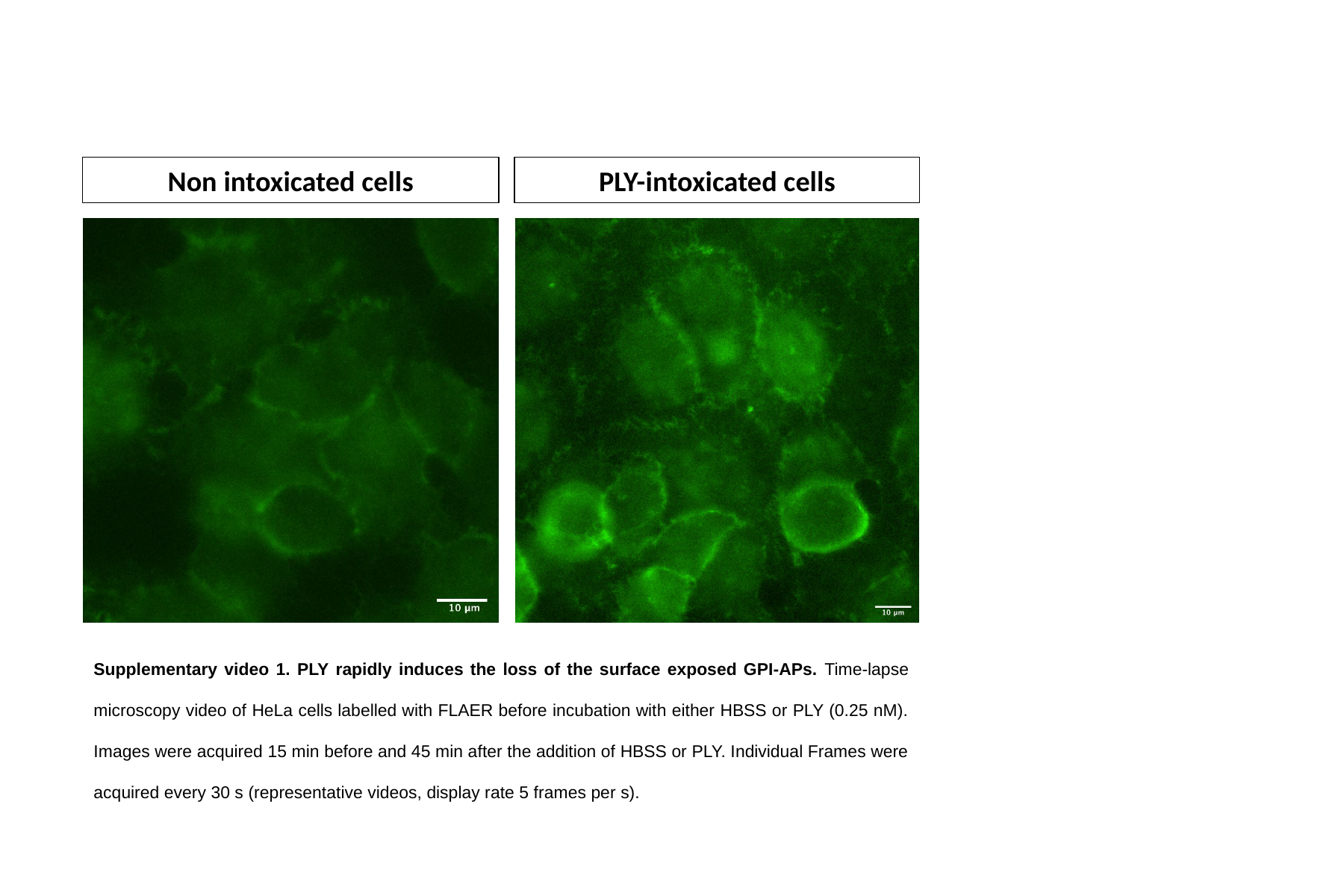

Non intoxicated cells
PLY-intoxicated cells
Supplementary video 1. PLY rapidly induces the loss of the surface exposed GPI-APs. Time-lapse microscopy video of HeLa cells labelled with FLAER before incubation with either HBSS or PLY (0.25 nM). Images were acquired 15 min before and 45 min after the addition of HBSS or PLY. Individual Frames were acquired every 30 s (representative videos, display rate 5 frames per s).

### Slide 2
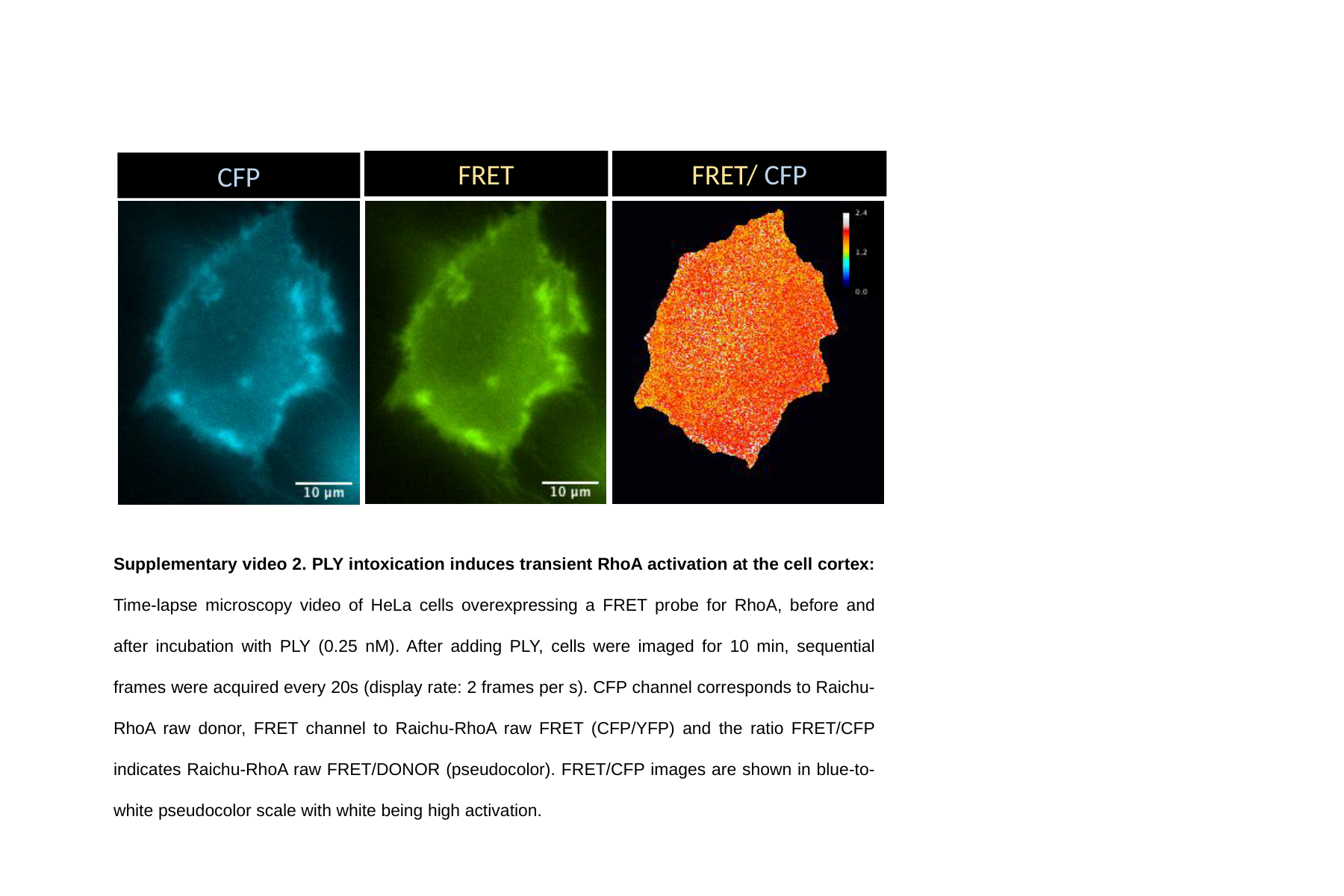

FRET
FRET/ CFP
CFP
Supplementary video 2. PLY intoxication induces transient RhoA activation at the cell cortex: Time-lapse microscopy video of HeLa cells overexpressing a FRET probe for RhoA, before and after incubation with PLY (0.25 nM). After adding PLY, cells were imaged for 10 min, sequential frames were acquired every 20s (display rate: 2 frames per s). CFP channel corresponds to Raichu-RhoA raw donor, FRET channel to Raichu-RhoA raw FRET (CFP/YFP) and the ratio FRET/CFP indicates Raichu-RhoA raw FRET/DONOR (pseudocolor). FRET/CFP images are shown in blue-to-white pseudocolor scale with white being high activation.

### Slide 3
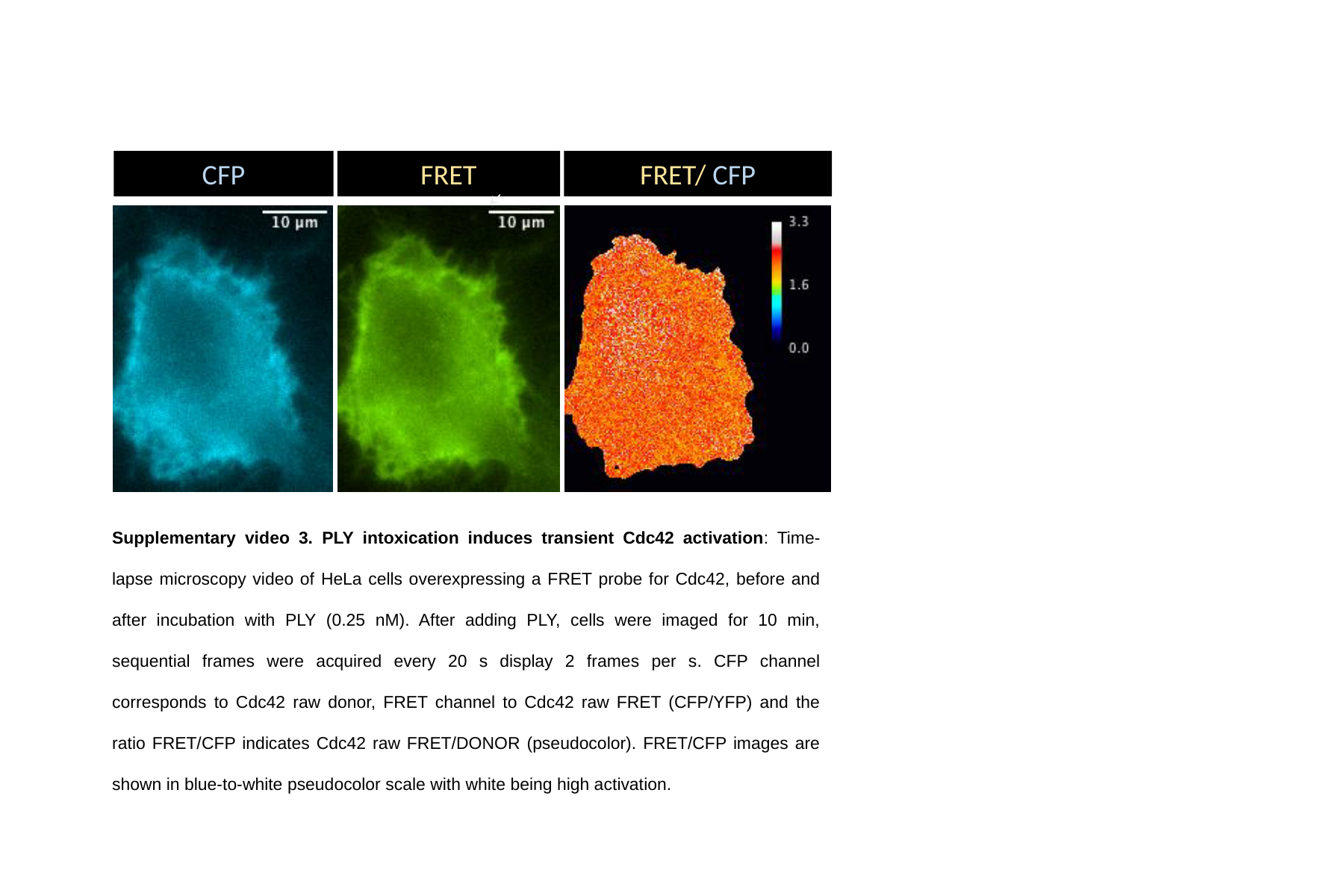

CFP
FRET
FRET/ CFP
Supplementary video 3. PLY intoxication induces transient Cdc42 activation: Time-lapse microscopy video of HeLa cells overexpressing a FRET probe for Cdc42, before and after incubation with PLY (0.25 nM). After adding PLY, cells were imaged for 10 min, sequential frames were acquired every 20 s display 2 frames per s. CFP channel corresponds to Cdc42 raw donor, FRET channel to Cdc42 raw FRET (CFP/YFP) and the ratio FRET/CFP indicates Cdc42 raw FRET/DONOR (pseudocolor). FRET/CFP images are shown in blue-to-white pseudocolor scale with white being high activation.
